# Targeted genome editing restores auditory function in adult mice with progressive hearing loss caused by a human microRNA mutation

**DOI:** 10.1101/2023.10.26.564008

**Authors:** Wenliang Zhu, Wan Du, Arun Prabhu Rameshbabu, Ariel Miura Armstrong, Stewart Silver, Yehree Kim, Wei Wei, Yilai Shu, Xuezhong Liu, Morag A Lewis, Karen P. Steel, Zheng-Yi Chen

**Author notes:** These authors contributed equally: Wenliang Zhu, Wan Du. Corresponding author. (Z.C.).

## Abstract

Mutations in *microRNA-96* (*MIR96*) cause dominant delayed onset hearing loss DFNA50 without treatment. Genome editing has shown efficacy in hearing recovery by intervention in neonatal mice, yet editing in the adult inner ear is necessary for clinical applications. Here, we developed an editing therapy for a C>A point mutation in the seed region of the *Mir96* gene, *Mir96^14C>A^* associated with hearing loss by screening gRNAs for genome editors and optimizing Cas9 and sgRNA scaffold for efficient and specific mutation editing in vitro. By AAV delivery in pre-symptomatic (3-week-old) and symptomatic (6-week-old) adult *Mir96^14C>A^* mutant mice, hair cell on-target editing significantly improved hearing long-term, with an efficacy inversely correlated with injection age. We achieved transient Cas9 expression without the evidence of AAV genomic integration to significantly reduce the safety concerns associated with editing. We developed an AAV-sgmiR96-master system capable of targeting all known human *MIR96* mutations. As mouse and human *MIR96* sequences share 100% homology, our approach and sgRNA selection for efficient and specific hair cell editing for long-term hearing recovery lays the foundation for future treatment of DFNA50 caused by *MIR96* mutations.

## INTRODUCTION

Hearing loss is a multi-factorial condition affecting a significant portion of the global population (World Health Organization, http://www.who.int/en/). Single genetic mutations causing hearing loss account for more than 50% of all congenital sensorineural hearing loss (SNHL)^1^, yet no pharmaceutical drug or biological treatment is available to slow down or reverse genetic deafness^2–4^. Among these deafness genes, microRNAs (miRNAs), which are short (∼20–24nt) endogenous non-coding RNAs that play a crucial role in the regulation of the expression of protein-coding genes, are considered important factors in the development of the inner ear and are required for normal hearing^5,6^. Specifically, mutations in *MIR96* (microRNA-96; miR-96) have been identified as causative factors for non-syndromic progressive hearing loss DFNA50 in humans and mice^7–11^. This mutation in the seed region of the human *MIR96* is the first example of point mutations in the mature sequence of a miRNA with an etiopathogenic role in a human Mendelian disease^6,11,12^.

*MIR96* is specifically expressed in the sensory cells of the inner ear^13^ and plays a critical role in cochlear development and the maintenance of hearing by reducing expression of the mRNAs of multiple target genes^8,14–16^. Point mutations in the seed region of *MIR96* lead to late onset progressive hearing loss with dominant inheritance patterns in human families^10,11^. Mice carrying an ENU-induced mutation, diminuendo (Dmdo), exhibit the phenotype of progressive hearing loss^17^. Homozygous Dmdo mice (*Mir96^Dmdo/Dmdo^*) and homozygous null mice (knockout of both *Mir96* and *Mir183*, *Mir96^dko/dko^*) are completely deaf, displaying abnormal hair cell stereocilia bundles and early-onset reduced hair cell numbers, demonstrating the requirement for *Mir96* in normal hair cell developmental^8,18^. In contrast, heterozygous *Mir96* null mice (*Mir96^/dko-^*) maintain normal hearing, while heterozygous Dmdo mice (*Mir96^Dmdo/+^*) develop adult-onset non-syndromic progressive hearing loss, suggesting that the hearing loss phenotype arises from the gain of novel target genes rather than loss of function^18^. In humans, heterozygous *MIR96* mutations also result in late onset non-syndromic progressive hearing loss, suggesting a similar gain of function due to the mutation. Delayed onset and progressive hearing loss offers a window of opportunity for intervention for hearing rescue.

In addition to its dominant inheritance pattern, overexpression of *Mir96* has been found to cause cochlear defects leading to hearing loss, rendering traditional gene therapy via ectopic *Mir96* expression unsuitable^9^. However, genome editing technologies that target the dominant *MIR96* mutations hold great promise to treat DFNA50. In vivo Cas9 nuclease delivery has demonstrated effectiveness in disrupting dominant alleles in multiple hearing loss disease models, including Lipofectamine RNP delivery^19,20^ or adeno-associated virus (AAV)-mediated delivery^21–26^. Inner ear base editing has also been employed to repair a *Tmc1* dominant mutation in mice^27^. In all the editing studies of genetic hearing loss, the interventions were done in neonatal mice with an immature cochlea^28^. In contrast, newborn human inner ears are fully developed^29^. The mouse cochlea undergoes significant developmental changes from the neonatal to adult stages, including changes in size, structure, and function^28^. This distinction highlights the need to evaluate the efficacy, safety, and time window of genome editing treatment in the mature cochlea to establish the feasibility of intervention for potential clinical applications.

In this study, we performed editing by NHEJ to target a dominant *Mir96* mutation (*Mir96^tm^*^3^*^.1Wtsi^*, referred to as *Mir96^14C>A^* in this report; equivalent of human rs587776523, g.7:129414596G>T in GRCh38) by AAV-mediated adult delivery in mice with hearing loss. We showed that AAV-mediated adult delivery of the editing complex was well-tolerated without impairing normal hearing. We screened sgRNAs for different editors and identified one for KKH-saCas9 for further modification, resulting in efficient and specific editing in vitro and in vivo. We tested interventions before and after the onset of hearing loss and established a time window when the treatment results in sustained and significant hearing preservation. To improve the safety profile of the editing strategy, we established the dosing without detectable AAV integrations and showed the lack of KKH-saCas9 expression 12 weeks post injection. As human and mouse *Mir96* share 100% sequences in the seed region, we developed a dual AAV system capable of targeting multiple human *MIR96* mutations. We successfully demonstrated the efficiency and specificity of this dual AAV system in *MIR96* mutant human cell lines, providing evidence for its potential generalization and broad utility. In conclusion, our study establishes the feasibility of in vivo genome editing as a viable approach for treating hearing loss caused by genetic mutations in adult animals.

## RESULTS

### A human *microRNA-96* mutation *Mir96^14C>A^* causes dominant adult-onset progressive hearing loss in mice

MiR-96 is highly expressed in mammalian sensory cells, including the inner ear, retina, and dorsal root ganglia, and functions as a master regulator controlling expression of many genes^8,17^. *Mir96* is super conserved across vertebrates, from zebrafish to humans, with the nucleotides within the *Mir96* seed region showing 100% identity in DNA sequence (Fig.1a).

**Fig. 1.**
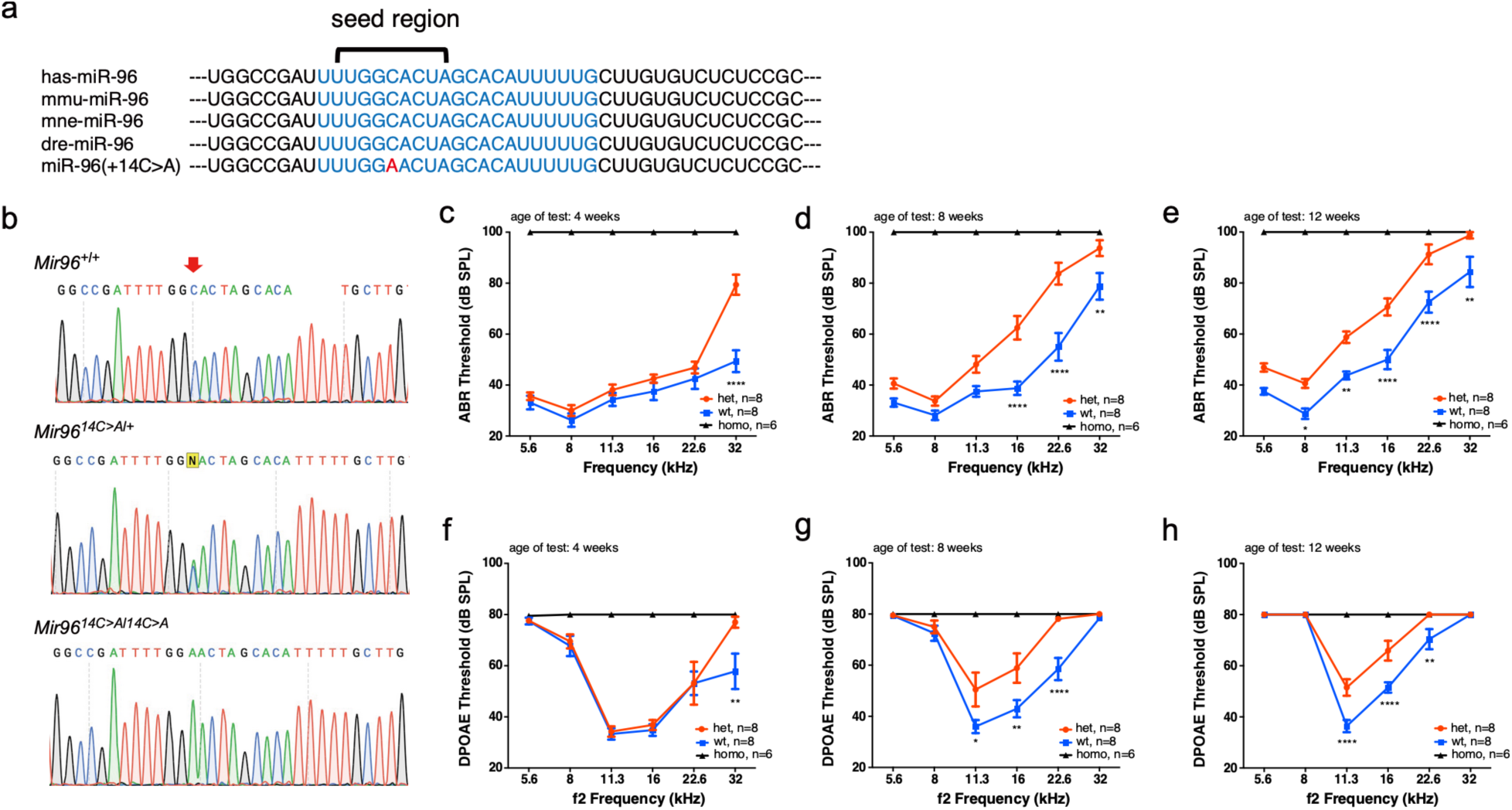
*Mir96^14C>A^*mutant mice present adult onset progressive hearing loss. **a**, *Mir96* sequence in human, mouse, macaque and zebrafish and 14C>A mutation. The blue text region shows 100% conservation. The mutation nucleotide in *Mir96^14C>A^*is displayed in red. **b**, Representative Sanger sequencing results showed the *Mir96* mutation locus in WT mice, *Mir96^14C>A/+^*, *Mir96^14C>A^/^14C>A^*. The red arrow indicates the mutated nucleotide. **c-e,** ABR thresholds in *Mir96^14C>A/+^*ears (red), compared to wild-type ears (blue) and *Mir96^14C>A/14C>A^* ears (black) at 4 weeks (c), 8 weeks (d), and 12 weeks (e), respectively. **f-h,** DPOAE thresholds in *Mir96^14C>A/+^*ears (red), compared to wild-type ears (blue) and *Mir96^14C>A/14C>A^* ears (black) at 4 weeks (f), 8 weeks (g), and 12 weeks (h), respectively. **i-n,** ABR thresholds in *Mir96^14C>A/+^*ears (red), compared to wild-type ears (blue) and *Mir96^14C>A/14C>A^* ears (black) at 5.6 kHz (i), 8 kHz (j), 11.3 kHz (k), 16 kHz (l), 22.6 kHz (m) and 32 kHz (n), respectively. Significant hearing loss, shown by the elevated ABR and DPOAE thresholds, was seen at 8 weeks, which became more severe at 12 weeks. N=8. Values and error bars reflect mean ± SD. Statistical analyses were performed by two-way ANOVA with Bonferroni correction for multiple comparisons: P value style, <0.05 (*), <0.01 (**), <0.001(***), <0.0001(****).

Two mutations in the seed region, +13G >A (rs587776522) and +14C >A (rs587776523), have been identified as the cause of autosomal dominant non-syndromic progressive hearing loss DFNA50^11^. Our mouse model carries the *Mir96^14C>A^*mutation, which substitutes the fourteenth nucleotide of cytosine in the conserved seed region with adenine^30^ (Fig.1a, b). To evaluate how hearing is affected by the mutation, we performed auditory brain stem response (ABR) and distortion product otoacoustic emission (DPOAE) tests on *Mir96^14C>A^* mice (Fig. 1c-h). *Mir96^14C>A/14C>A^* mice exhibited complete hearing loss across all frequencies and age groups by ABR and DPOAE at 4 weeks of age (Fig. 1c, f). *Mir96^14C>A/+^*mice at 4 weeks of age, we observed a 30 dB elevation in ABR and a 19 dB elevation in DPOAE at the frequency of 32 kHz but no significant changes in ABR/DPOAE thresholds at other frequencies compared to wild-type (WT) ears (Fig. 1c, f), indicating the onset of hearing loss starting at 4 weeks at 32 kHz. By 8 weeks, the ABR thresholds were significantly elevated from 16 to 32 kHz, with an average increase of 20 dB in *Mir96^14C>A/+^* ears compared to WT ears. Similarly, DPOAE thresholds were significantly elevated by an average of 16 dB from 11.3 to 22.6 kHz frequencies *Mir96^14C>A/+^* ears compared to WT ears (Fig. 1g). At 12 weeks, the hearing loss became more severe, as ABR thresholds were significantly elevated across all frequencies when compared to WT ears, with an average elevation of 15 dB. Additionally, DPOAE thresholds were significantly elevated 15 dB on average compared to WT ears, particularly at 11.3 and 16 kHz (Fig. 1h). These results showed that *Mir96^14C>A/+^* mice exhibit adult-onset hearing loss starting at 4 weeks at the high frequency of 32kHz, which becomes progressive with further elevation of ABR/DPOAE thresholds at 8 and 12 weeks across other frequencies.

### Screening of genome editing systems in mouse and human cells

To specifically disrupt the *Mir96^14C>A^* allele, sgRNAs were designed to target the allele for each of five types of CRISPR nucleases: spCas9^31^, scCas9++^32^, KKH-saCas9^33^, sauriCas9^34^, LZ3 Cas9^35^ (Fig. 2a). All Cas9/sgRNA combinations were in the same pMAX-Cas9/sgRNA backbone with three nuclear localization signals (NLSs) to ensure consistency in plasmid structure (Fig. 2b and Extended Data Fig. 1a). Using primary *Mir96^14C>A/+^* fibroblasts, spCas9/sgRNA-1 exhibited the highest editing efficiency (14.3± 2.3%) (Fig. 2c, d). LZ3 Cas9/sgRNA-1 and KKH-saCas9/sgRNA-4 also showed comparable editing efficiency (12.7± 2.0% and 10.1± 2.1%), while scCas9++/sgRNA-1, scCas9++/sgRNA-2 and sauriCas9/sgRNA-4 displayed modest indel frequency (Fig. 2c, d). spCas9/sgRNA-1 and KKH-saCas9/sgRNA-4 exhibited high specificity in targeting the *Mir96^14C>A^*allele, shown by next generation sequencing (NGS) analysis that all the indels occurred exclusively at the mutant allele and were undetectable in *Mir96*^+/+^ cells (less than 0.03%) (Fig. 2d-f and Extended Data Fig. 1b). We compared the editing efficiency of Cas9/sgRNA ribonucleoprotein (RNP) and plasmids in primary fibroblasts of *Mir96^14C>A/+^*mice. RNP nucleofection yielded the highest editing efficiency, while lipofectamine-mediated RNP delivery was less efficient (Extended Data Fig. 1c). The AAV systems with spCas9/sgRNA1 dual AAV plasmids (7.3± 1.5%) and KKH-saCas9/sgRNA4 (10.4± 1.8%) single AAV plasmid exhibited editing frequencies comparable to the pMAX-Cas9 plasmid transfection (14.3± 2.3%) (Extended Data Fig. 1c).

**Fig. 2.**
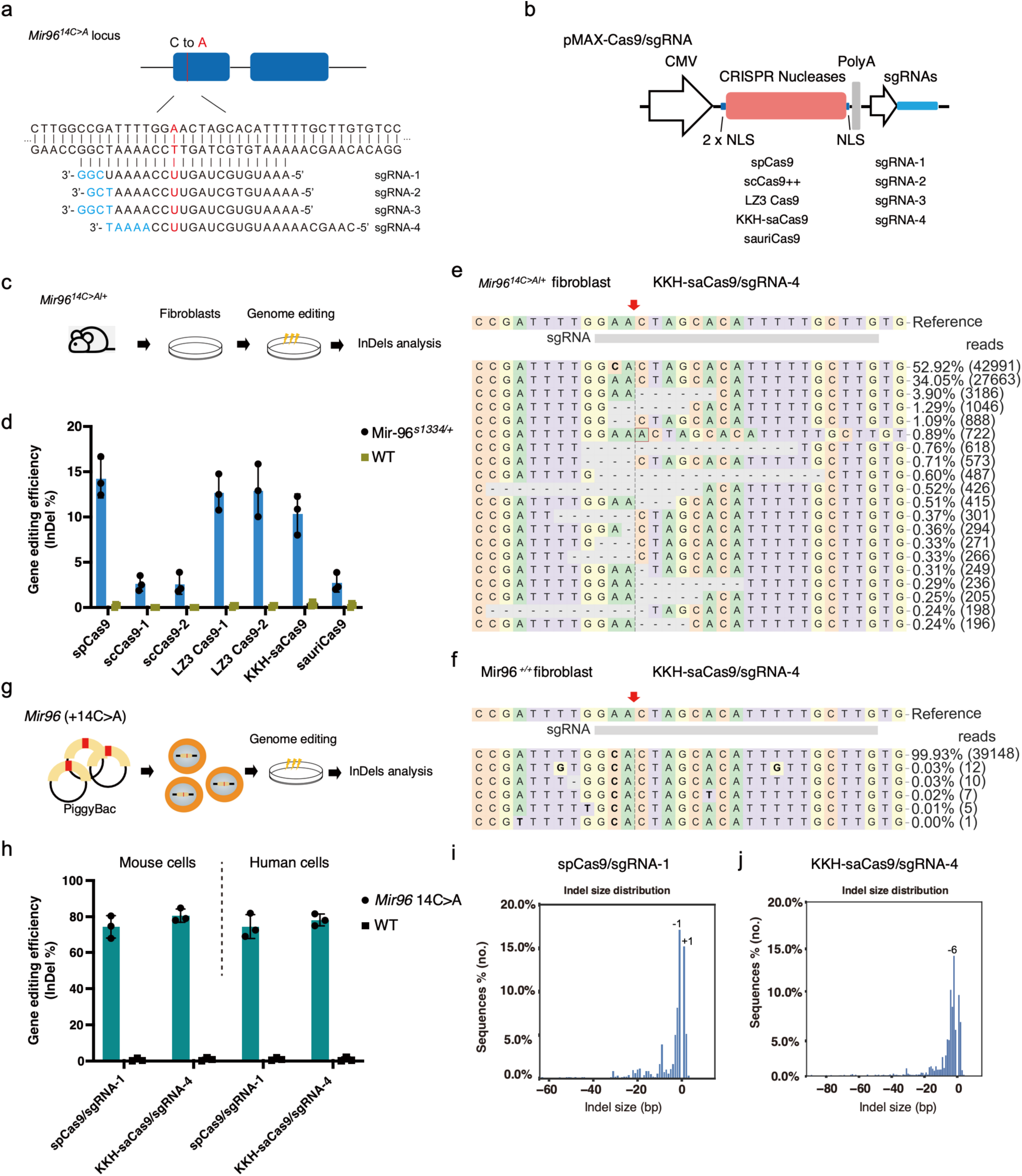
*Mir96 14C>A* allele specific genome editing using different CRISPR systems in mouse and human cells. **a**, Sequence of the *Mir96* +14 C>A mutation loci and the sgRNAs designs. The mutated nucleotide in the *Mir96^14C>A^* allele is displayed in red. The protospacer adjacent motifs (PAMs) nucleotides are displayed in blue. **b**, Schematic overview of plasmid constructions for different CRISPR systems used for in vitro screening. **c**, Experiments design of preparation and genome editing of primary fibroblasts from *Mir96^14C>A/+^* mice. **d**, Bar chart of the indel frequency in *Mir96^14C>A/+^* and wild-type primary fibroblast after genome editing using different Cas9/sgRNA combinations. n=3. Each dot represents a unique sequencing reaction. Values and error bars reflect mean ± SD. **e, f**, Representative NGS results from KKH-saCas9/sgRNA-4 edited *Mir96^14C>A/+^* and wild-type primary fibroblasts. The red arrow indicates the double-stranded DNA cutting site. **g**, Schematic overview of establishment of human *MIR96* +14 C>A HEK-293T cell line and mouse *Mir96^14C>A^* NIH-3T3 cell line using PiggyBac system and genome editing procedure. **h**, Bar chart showing the editing efficiency of *Mir96* mutation locus and wild-type locus using spCas9/sgRNA-1 and KKH-saCas9/sgRNA-4 in human *MIR96* +14 C>A HEK-293T cells and mice *Mir96* +14 C>A NIH-3T3 cells. n=3. Each dot represents a unique sequencing reaction. Values and error bars reflect mean ± SD. **i, j**, Indel profiles from spCas9/sgRNA-1 and KKH-saCas9/sgRNA-4 edited *Mir96* +14C>A HEK-293T cells. Minus numbers represent deletions, plus numbers represent insertions.

Due to 100% homology in the *Mir96* seed region across species, the sgRNAs design targeting the *Mir96^14C>A^* allele in mice can be used in the human *MIR96* with the same mutation. To validate the editing systems in human cells, we generated human and mouse cell lines carrying the *Mir96^14C>A^* mutation using the PiggyBac transposon system (Fig. 2g). In cells transfected with spCas9/sgRNA-1 and KKH-saCas9/sgRNA-4, NGS analysis showed a high level of indel formation in both human and mouse *Mir96* +14C>A cell lines, where negligible indels were observed in the *Mir96* wild-type cell lines (Fig. 2h). The indel profile showed various types of indels for spCas9/sgRNA-1 and KKH-saCas9/sgRNA-4, with single nucleotide deletions (-1 and +1) being the most common type for spCas9/sgRNA-1, and a six-nucleotide deletion being the most common for KKH-saCas9/sgRNA-4 (Fig. 2i, j).

In addition to disrupting the mutated allele with CRISPR nucleases, we also attempted to correct the *Mir96^14C>A^*mutation using prime editing^36–38^(Extended Data Fig. 2a). We designed a pegRNA located upstream of the mutate nucleotide, containing pegRNA-extensions with 13 bp primer binding sites (PBSs) and 16 bp RT-templates harboring the corrected sequence (Extended Data Fig. 2b). We transfected HEI-OC1 cells carrying the *Mir96* +14C>A mutation with plasmids encoding primer editors and pegRNA. NGS analysis showed that prime editing successfully corrected the mutation in *Mir96^14C>A^* cells with an efficiency of 2.8±1.4% (Extended Data Fig. 2c, d). However, due to the modest efficiency of prime editing at this locus and the lack of an efficient in vivo delivery system for the inner ear, CRISPR nuclease-mediated knock-out of the mutated allele is a more suitable choice for treating hearing loss caused by the *Mir96* mutation.

### Optimization of CRISPR/sgRNA delivery system for mature cochlea in adult mice

We chose AAV serotype 2 (AAV2), which has demonstrated high efficiency and specificity in transducing both mature inner hair cells (IHCs) and outer hair cells (OHCs) compared to other AAV serotypes^39,40^, making AAV2 an excellent candidate vehicle for delivering Cas9/sgRNA to hair cells. To further assess the delivery efficiency of AAV2, we initially delivered AAV2 carrying a GFP cassette into the adult cochlea through round window membrane and canal fenestration (RWM+CF) injection in 18-week-old mice (Extended Data Fig. 3a). Four weeks after injection, robust GFP expression was observed in nearly all IHCs throughout the cochlear turns and robust OHC transduction with an apex-to-base gradient (Extended Data Fig. 3a-d). Importantly, the majority of GFP-positive cells were hair cells, with only limited GFP+ cells observed in other cell types, including supporting cells, indicating that AAV2 specifically and efficiently targets auditory hair cells in adult mice.

To maximize editing efficiency in vivo to improve hearing rescue, we modified AAV Cas9/sgRNA construction by incorporating a bipartite nuclear localization signal (bpNLS), an sv40 NLS at the N-termini, and another bpNLS at the C-termini (Fig. 3a) to enhance nuclear import and their access to genomic DNA^41,42^. The KKH-saCas9 sgRNAs were optimized by using an 84 nt sgRNA (crRNA-tracrRNA duplex extension) which exhibits higher activity than the canonical full length sgRNA^33^. Additionally, we introduced a 3rd A-U flip in the stem loop to remove a putative RNA Pol III terminator sequence (4 consecutive U’s), further enhancing expression levels under the U6 promoter^43^ (Fig. 3b, c). The optimized construct was tested in the primary fibroblasts derived from Ai14 mice, which contain “CAG-STOP-tdTomato” cassette^44^. In the cells transfected with the optimized KKH-saCas9/sgRNA constructs, an increase in editing efficiency of approximately 3.05-fold and 1.98-fold was detected for KKH-saCas9/sgtdT-1 and KKH-saCas9/sgtdT-2 compared to the original designs, respectively, shown by the tdT^+^ cells (Fig. 3d, e). The editing efficiency was also evaluated in HEK-GFP cells using KKH-Cas9/sgGFP targeting GFP, and the ratio of GFP-negative cells showed significantly higher efficiency when using the optimized AAV constructions compared to the original design (Extended Data Fig. 3e, f). The optimized AAV plasmid design and gRNA modifications were used for the production of AAV-KKH-saCas9/sgRNA-4 for the in vivo editing study.

**Fig. 3.**
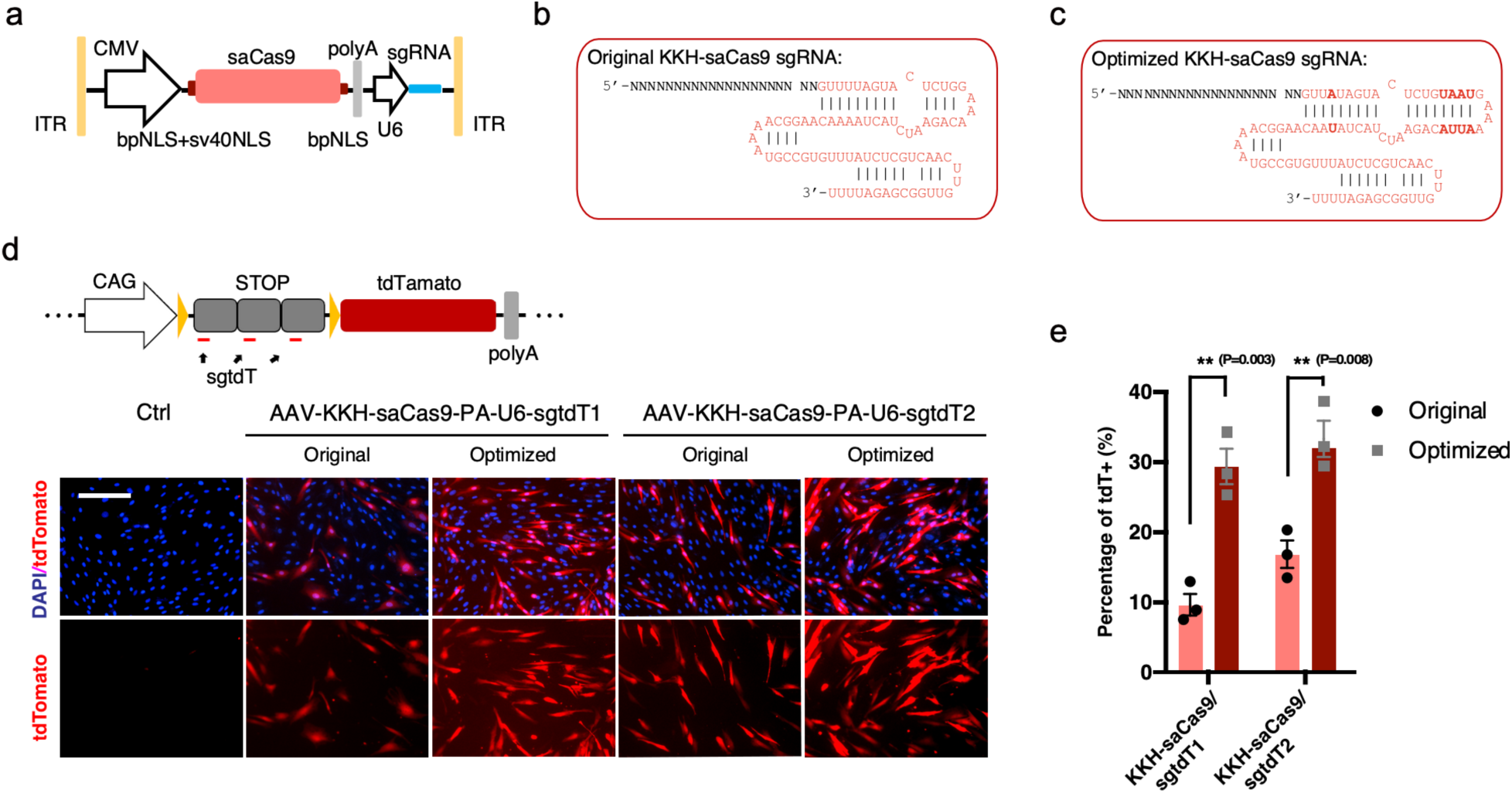
Optimization of CRISPR constructs for inner ear delivery. **a**, The design of the optimized AAV structure of KKH-saCas9/sgRNA vector with multiple NLS sites. **b, c**, Sequence of unmodified and optimized KKH-saCas9 sgRNA, with sequence changes in bold. **d**, Schematic view and representative fluorescence images of tdTomato reporter from primary fibroblasts after editing by unmodified and optimized KKH-saCas9/sgRNA systems. tdTomato^+^ cells are edited. **e**, Bar chart showing the editing efficiency by the quantification of tdTomato^+^ cells after editing. Values and error bars reflect mean ± SD. Each dot represents one independent experiment.

### In vivo genome editing of *Mir96^14C>A^* allele in mature cochlea

KKH-saCas9/sgRNA-4 was packaged into AAV2 (Fig. 4a) and delivered to adult *Mir96^14C>A/+^* cochlea (6 weeks old) via round window membrane and canal fenestration (RWM+CF) injection at a dose of 6×10^9^ vg (Fig. 4a). Cochleae were collected 4- and 8-weeks post-injection for NGS (Fig. 4b). In vivo NGS of cochlear tissue detected the presence of indels at the *Mir96* mutant locus in the injected ear at 4 and 8 weeks, while no indels were detected in the contralateral uninjected ears (Fig. 4c, d and Extended Data Fig. 4a-c). Importantly, AAV2-KHH-SaCas9-sgRNA-4 specifically edited the *Mir96^14C>A^* allele, as no indels were observed in AAV2-KKH-saCas9/sgRNA-4 treated wild-type (WT) mice or AAV2-KKH-saCas9/sgCtrl treated *Mir96^14C>A/+^* mice (Fig. 4d and Extended Data Fig. 4a-c). After 8 weeks of injection, the indel rate was determined to be 0.76 ± 0.13%, slightly higher than the rate observed 1 month after injection (0.63 ± 0.14%) (Fig. 4d). However, the indel frequency from organ of Corti samples does not accurately reflect the editing efficiency in hair cells, which constitutes less than 5% of cochlear cells.

**Fig. 4.**
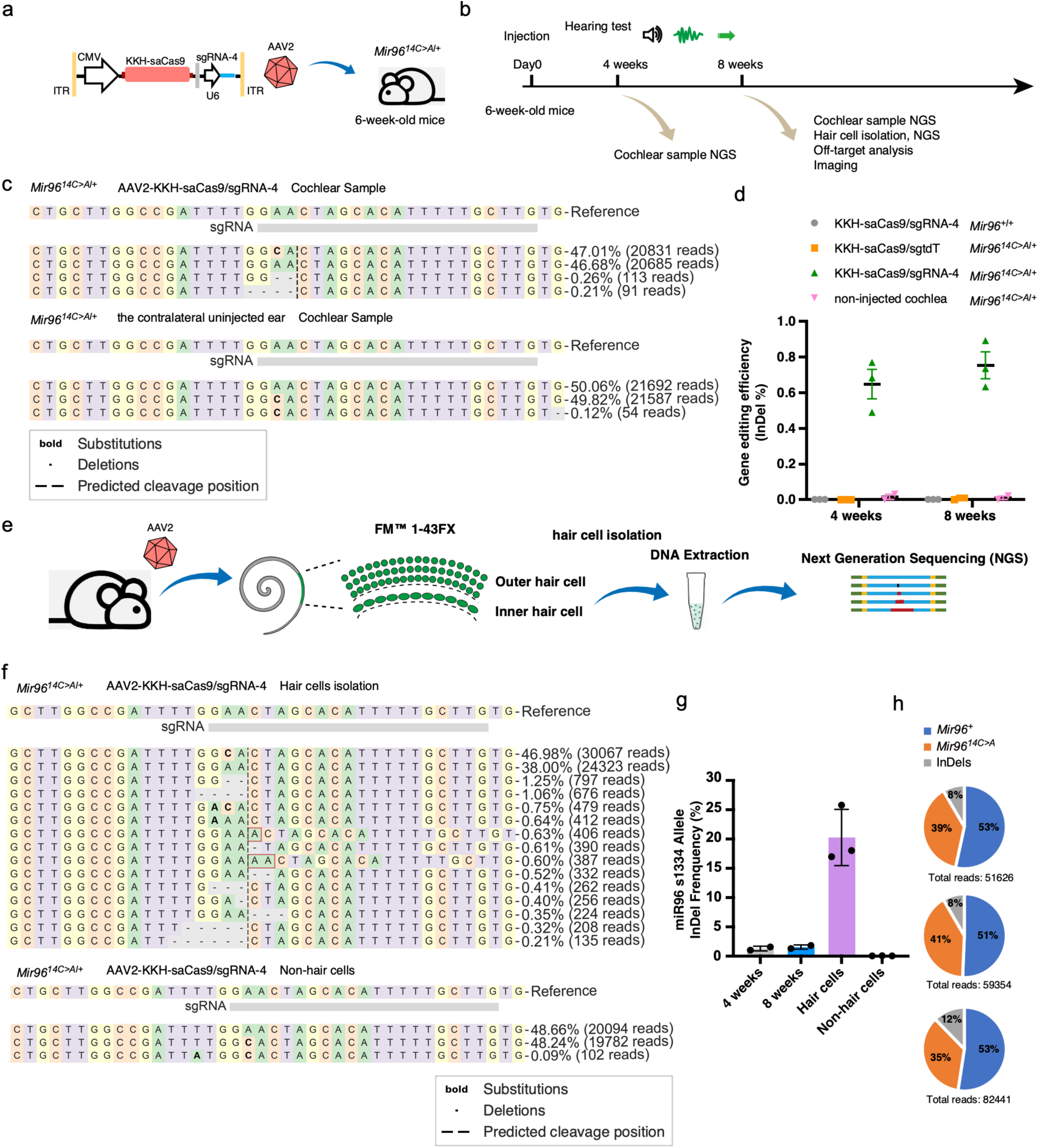
AAV2-CRISPR mediated targeted genome editing at *Mir96^14C>A^* locus in hair cells of adult *Mir96^14C>A/+^* mice. **a**, Schematic view of the construction and packaging of AAV2-CMV-KKH-saCas9-sgRNA4 for genome editing in adult mice. **b**, Experimental overview for in vivo studies. AAV2 was injected in the inner ear of 6-week-old mice, and one group was used for sequencing 4 and 8 weeks later. **c**, Representative NGS results of cochlea samples from AAV2-KKH-saCas9-sgRNA-4 injected and the contralateral uninjected ears of *Mir96^14C>A/+^* mice. **d**, Quantification of indel frequency of the NGS results from AAV2-KKH-saCas9-sgRNA-4 injected, AAV2-KKH-saCas9-sgCtrl injected and uninjected ears from *Mir96^14C>A/+^* mice, as well as AAV2-KKH-saCas9-sgRNA-4 injected ears from wild-type mice (n=9). Cochleae were collected 4 weeks and 8 weeks after AAV injection. In uninjected mice, background indel frequencies ranged between 0% and 0.05%. Each dot represents a unique sequencing reaction from a combination of 3 cochleae. Values and error bars reflect mean ± SD. **e**, Schematic overview of the experimental protocol of hair cell isolation, cell lysis and NGS. **f**, Representative NGS result of isolated hair cells from AAV2-KKH-saCas9-sgRNA-4 injected cochlea. **g**, Quantification of *Mir96^14C>A^*allele-specific indel frequency from NGS of hair cell and cochlea samples after AAV2-KKH-saCas9-sgRNA-4 injection (n=3 for each). Each dot represents a unique sequencing reaction from 3 cochleae combination. Values and error bars reflect mean ± SD. **h**, Percentage of *Mir96* wild-type allele reads, 14C>A reads, and indel-containing reads in the NGS results from AAV2-KKH-saCas9-sgRNA-4 injected hair cells from 3 independent experiments.

To obtain a precise measurement of editing efficiency in hair cells, we performed injection and assessed isolated hair cells after labeling hair cells with the FM1-43 uptake assay for NGS assay. FM1-43 and its fixable analog FM1-43FX can specifically label hair cells by passing through open mechanotransduction channels^45,46^ (Extended Data Fig. 4d). The labeled cells were hand-picked under an inverted fluorescent microscope, lysed, with DNA extracted for PCR and NGS to study indels (Fig. 4e). Indel-containing *Mir96* sequencing reads from injected *Mir96^14C>A^*^/+^ mice allowed us to directly assess the allele specificity of *Mir96^14C>A^* allele in vivo. Robust indel formation was observed in isolated hair cells from injected animals 8 weeks after treatment, with an indel frequency of 16.3-25.73% in the *Mir96^14C>A^*allele and a wider range of indel types including -2bp, -4bp, +1bp, -1bp, compared to the indels observed in cochlear samples (Fig. 4f, g), indicating efficient targeted genome editing in hair cells. From three independent experiments, the ratio of *Mir96* wild-type allele reads to *Mir96^14C>A^* allele reads was analyzed based on NGS results after editing. In the injected ear, the ratio between the *Mir96^14C>A^* allele that combined unedited with indel-containing reads and that of *Mir96* wild-type reads was 52.3%:47.7% (Fig. 4h), similar to that observed in uninjected ears (Extended Data Fig. 4e), indicating no noticeable chromatin lesion, insertion or large deletions caused by genome editing.

### In vivo genome editing improved short- and long-term hearing preservation

We injected AAV2-KKH-saCas9-sgRNA-4 into the cochlea of 6-week-old (P40-P45) *Mir96^14C>A/+^* mice through the round window membrane (RWM) with canal fenestration, with the contralateral uninjected ears serving as controls. 10 weeks after the injection, at 16 weeks of age, the injected ears showed overall lower auditory brainstem response (ABR) thresholds compared to the uninjected control ears, at frequencies from 5.6 to 16 kHz (Fig. 5b). The ABR threshold reduction ranged from 12 dB at 5.6 kHz to 18 dB at 16 kHz for frequencies below 22.6 kHz, with an average reduction of 13 dB. Additionally, lower distortion product otoacoustic emission (DPOAE) thresholds were observed in the injected ears compared to the control ears at frequencies of 11.3 and 16 kHz, with reductions of 14 dB to 21 dB, respectively (Fig. 5c).

**Fig. 5.**
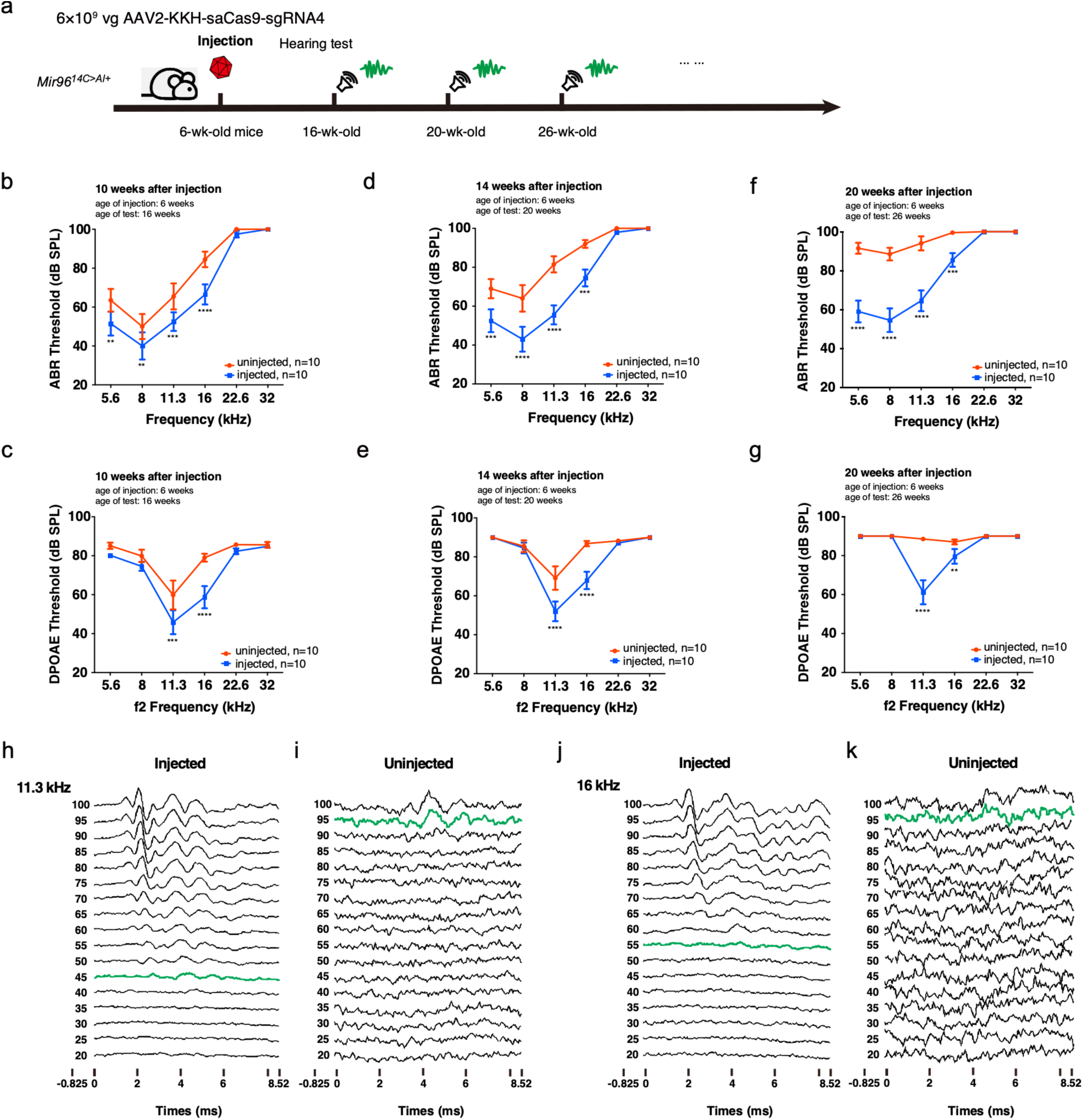
Sustained hearing rescue by AAV2-KKH-saCas9-sgRNA-4 in adult *Mir96^14C>A/+^* mice. **a.** Schematic view of workflow of AAV2-KKH-saCas9-sgRNA-4 constructions for auditory function assay with the editing complex delivered into adult *Mir96^14C>A/+^* mouse cochleae. **b-g.** After AAV2-CMV-KKH-saCas9-sgRNA-4 injection into 6-week-old *Mir96^14C>A/+^* ears, significant reductions in ABR (b, d, f) and DPOAE (c, e, g) thresholds were detected in injected (blue) vs. uninjected contralateral ears (red), at 16 weeks age **(b-c)**, 20 weeks age **(d-e)**, and 26 weeks age **(f-g)**. n=10. **h-k**. Representative ABR waveforms recorded from an injected (left) and an uninjected ear (right) of a mouse of 20 weeks of age at the frequencies of 11.3 kHz **(h-i)** and 16 kHz **(j-k)**. The thresholds were determined by the detection of peak 1 (green color traces). Values and error bars reflect mean ± SEM. Statistical analyses were performed by two-way ANOVA with Bonferroni correction for multiple comparisons: P value style, <0.05 (*), <0.01 (**), <0.001(***), <0.0001(****).

To assess long-term auditory function following hair cell-targeted genome editing, hearing tests were conducted over an extended period. 14 weeks after injection (i.e., 20 weeks of age), significantly lower ABR thresholds were detected in injected ears compared to the uninjected control ears at frequencies of 5.6, 8, 11.3, and 16 kHz, with an average reduction of 21 dB in ABR thresholds. At the frequencies of 8 and 11.3 kHz, the reduction is 21 and 26 dB, respectively (Fig. 5d). Significant reductions in DPOAE thresholds were observed in the injected ears compared to the uninjected control ears at the frequencies of 11.3 and 16 kHz, with reductions at 15 dB and 19 dB, respectively (Fig. 5e). 20 weeks post injection (26 weeks age), more dramatic lower ABR thresholds were detected in the injected ears compared to the uninjected control ears from 5.6 to 16 kHz, with an average reduction of 28 dB in ABT thresholds (Fig. 5f). For the frequencies of 5.6 and 8 kHz, the ABR thresholds were reduced by 33 dB and 34 dB, respectively. Similarly, dramatic lower DPOAE thresholds were detected in injected ears compared with uninjected control ears at frequencies of 11.3 and 16 kHz, with reductions of 27 dB and 7 dB at respectively (Fig. 5g). Representative ABR waveforms recorded from an AAV2-KKH-saCas9-sgRNA-4 injected ear (Fig. 5h-j) and the contralateral uninjected ear (Fig. 5i-k) 14 weeks post injection using 11.3 kHz (Fig. 5h-i) and 16 kHz (Fir. 5j-k) tone bursts at incrementally increasing sound pressure levels (SPLs) from 20 to 100 dB (Fig. 5h-k) further illustrated the improved hearing preservation. Collectively, these results demonstrate that AAV2-mediated genome editing effectively abolished the expression of the mutant *Mir96*, leading to improved short- and long-term hearing preservation.

Delayed onset progressive hearing loss caused by *MIR96* mutations offers a window of opportunity for intervention. To determine the optimal time window for genome editing treatment in *Mir96^14C>A/+^* mice, we further performed AAV2-KKH-saCas9-sgRNA-4 injections in 3-week-old mice in which initial hearing loss was detected only at the high frequency and in 16-week-old mice in which severe hearing loss was detected in all frequencies (Fig. 1c, f; Extended Data Fig. 5a). For 3-week-old injection, significantly lower ABR thresholds were detected 13 weeks post injection in AAV2-KKH-saCas9-sgRNA-4 treated ears compared to the untreated control ears, with an average ABR threshold reduction of 19 dB from 5.6 to 16 kHz (Extended Data Fig. 5b). A lower ABR threshold of 8 dB was seen at 22.6 kHz although the reduction is not statistically significant. The average ABR threshold reduction in 3-week-old *Mir96^14C>A/+^* mouse injection was greater than that in 6-week-old *Mir96^14C>A/+^* mouse injection, 19 dB vs. 13 dB. Significantly lower DPOAE thresholds were also detected in the AAV2-KKH-saCas9-sgRNA-4 treated ears compared to the contralateral control ears at frequencies of 11.3 and 16 kHz, with a reduction of 31 dB at 11.3 kHz and 24 dB at 16 kHz. Lower DPOAE threshold of 9 dB was detected at 22.6 kHz; however, it is not statistically different (Extended Data Fig. 5c). Again, the DPOAE threshold reduction in *Mir96^14C>A/+^* mice injected at 3-week-old was greater than that in *Mir96^14C>A/+^* mice injected at 6-week-old, 27 dB vs. 18 dB.

In the 16-week-old mice injection group, no difference in ABR and DPOAE thresholds was detected between AAV2-KKH-saCas9-sgRNA-4 treated and contralateral control ears at 26 weeks age (Extended Data Fig. 5d-e). The data demonstrate that at the later stage with severe hearing loss deterioration, genome editing therapy is no longer an effective treatment. Combined with the data from 3-, 6- and 16-week injections, the results highlight the importance of early intervention for more effective outcomes before substantial hearing loss has initiated.

### In vivo genome editing preserves hair cell survival

To test the effect of editing on HCs in vivo, we injected the AAV2-KKH-saCas9-sgRNA-4 into the inner ear of 6-week-old *Mir96^14C>A/+^* mice. Cochleae were harvested 10 weeks after the injection for hair cell labeling using an anti-MYO7A antibody and confocal imaging (Fig. 6a). In the uninjected *Mir96^14C>A/+^* inner ears, outer hair cell (OHC) loss across cochlear turns was observed with the most severe loss in the basal turn (Fig. 6c, e, f). A slight loss of the inner hair cells (IHC) was detected in the basal turn only (Fig. 6c, e, f). In the injected *Mir96^14C>A/+^* ears, improved OHC survival was seen across all frequency regions compared to the control group (Fig. 6b, d, f); whereas a slight reduction in the IHC number was seen in the base similar to uninjected ears (Fig. 6g). To further examine hair cell structure, scanning electron microscopy (SEM) was performed to visualize the details of OHC and IHC stereocilia. Cochleae from uninjected and AAV2-KKH-saCas9-sgRNA-4 injected *Mir96^14C>A/+^* mice were harvested at 14 weeks after injection and imaged by SEM (Fig. 6a). Hair cells from the uninjected *Mir96^14C>A/+^* mice cochleae showed signs of degeneration, including missing stereocilia in the outer hair cells and disorganized stereocilia in the inner hair cells (Fig. 6i, k). In contrast, the hair cells of injected *Mir96^14C>A/+^* mouse cochlea had organized and well-preserved stereocilia (Fig. 6h, j). These findings demonstrate that in vivo genome editing by AAV2-KKH-saCas9-sgRNA-4 rescues hair cells in *Mir96^14C>A/+^* mice by promoting their survival and maintaining the structure of stereocilia.

**Fig. 6.**
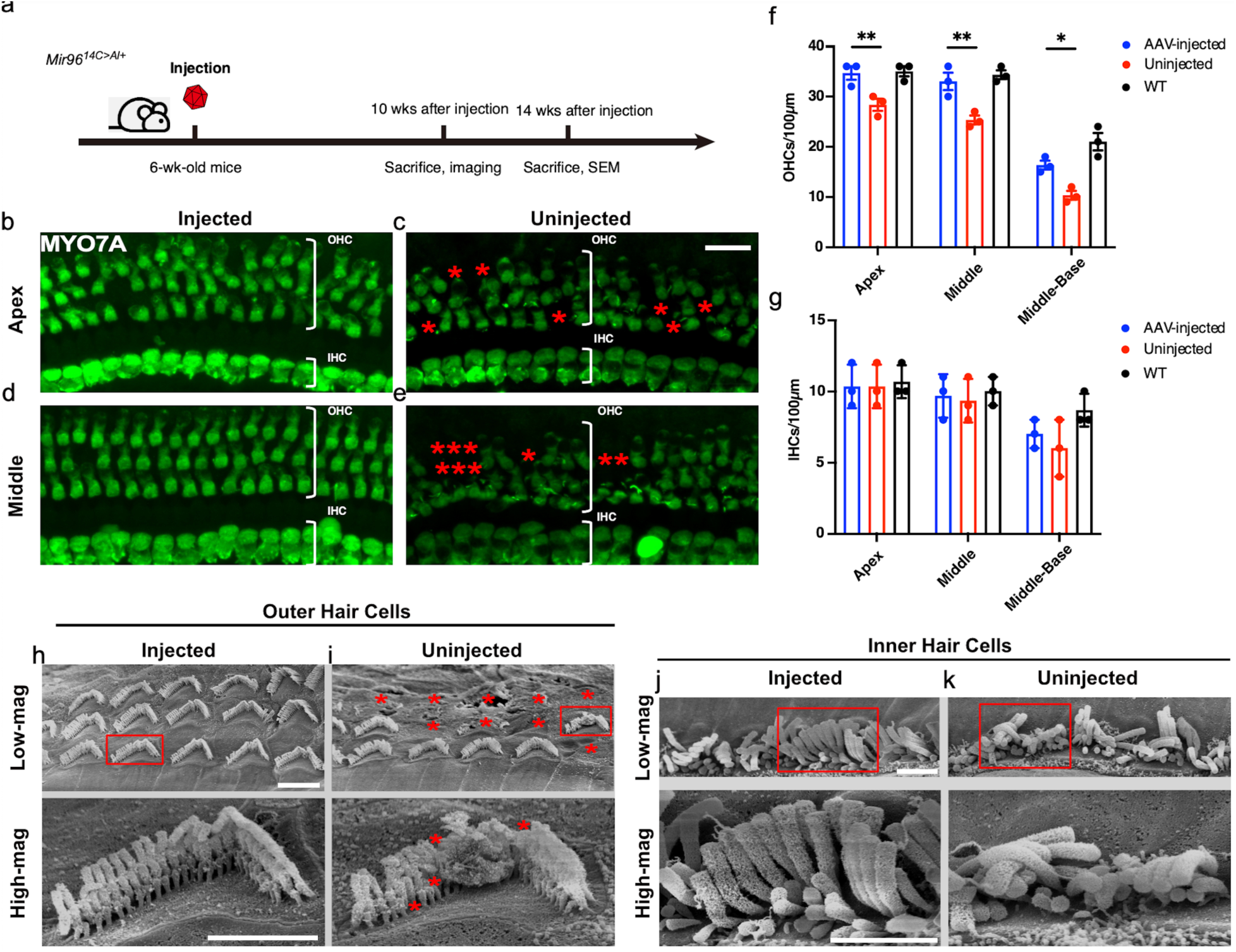
AAV2-KKH-saCas9-sgRNA-4 editing on hair cell survival and maintains stereocilia intergity in *Mir96^14C>A/+^* mice. **a,** Schematic diagram of experimental design, with confocal analysis and SEM study of the ears harvested 10 weeks and 14 weeks post injection, respectively. **b-e,** Representative confocal z-stack images of whole mount cochleae from uninjected (c, e) and AAV2-KKH-saCas9-sgRNA-4 injected (b, d) *Mir96^14C>A/+^* mice. Hair cells were stained for MYO7A (green). Asterisks point to missing hair cells (**i**). Scale bar, 20μm. **f, g,** Quantification and comparison of the number of OHCs (f) and IHCs (g) across the cochlear turns from injected and uninjected *Mir96^14C>A/+^* mice 10 weeks post injection. Error bar represents SD. **h, i,** Images of scanning electron microscopy (SEM) of injected (h) uninjected (i) *Mir96^14C>A/+^* outer hair cell bundles at the apical turn. The asterisks indicate the missing stereocilia in an outer hair cell from an uninjected ear. **j, k,** Images of SEM of injected (j) and uninjected (k) *Mir96^14C>A/+^* inner hair cell bundles at the apex turn. Scale bar, 2µm.

### Safety assessment of AAV-mediated genome editing treatment in cochleae of adult mice

Development of editing therapy for hearing loss requires safety assessment. To address potential safety concerns about AAV-delivered CRISPR genome editing in vivo, the chronic expression of saCas9 and the risk of AAV vector integration were evaluated. RT-PCR analysis revealed that KKH-saCas9 RNA levels peaked at 4 weeks, decreased at 8 weeks, and became undetectable after 12 weeks in the injected ear (Fig. 7a). Only trace levels of KKH-saCas9 mRNA were detected in the contralateral uninjected ear at 4 weeks post-injection (Fig. 7a). By Western blotting, we confirmed the presence of KKH-saCas9 protein at 4 weeks post-injection and its absence after 12 weeks (Fig. 7b). These findings collectively show that the expression of KKH-saCas9 expression was effectively shut off within three months after injection, likely due to the silencing of the CMV promoter, which is known to undergo silencing over time in vivo^47–49^.

**Fig. 7.**
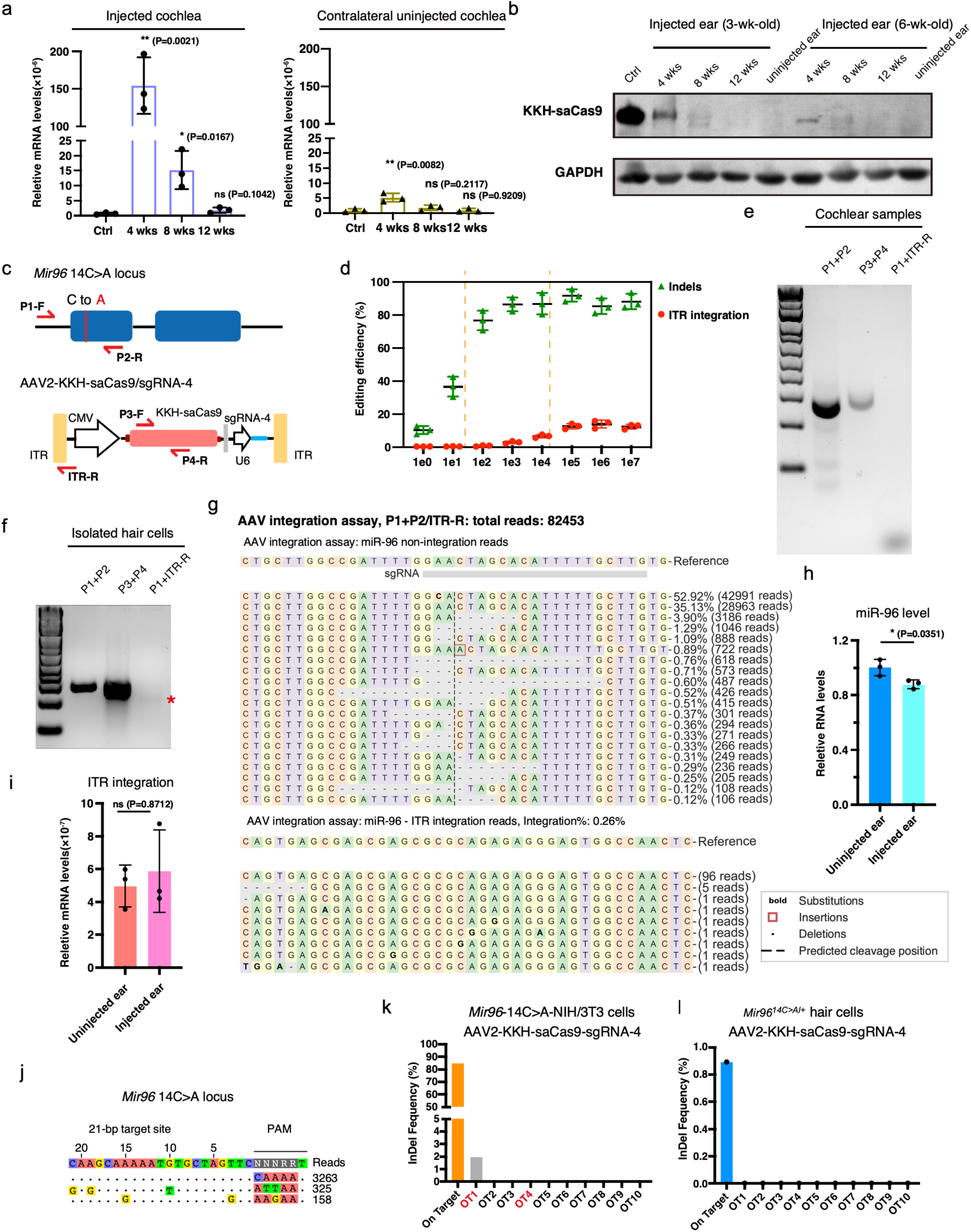
Safety assessment of AAV delivery of genome complex in adult mice. **a**, qPCR analysis of KKH-saCas9 mRNA level in the injected cochlea and the contralateral uninjected cochlea (n=6). Each dot represents an independent result from 2 cochleae combined Values and error bars reflect mean ± SD. **b**, Western blotting of KKH-saCas9 protein in injected cochlea and the contralateral uninjected cochlea, showing KKH-saCas9 protein level. **c**, Schematic diagram of the primers and design of AAV vector integration assay. **d**, Quantification of indel frequency by NGS results and AAV vector ITR integration ratio from AAV2-KKH-saCas9-sgRNA-4 edited cells. HEK-293T-miR96-14C>A cells were treated with different doses of AAV for 7 days before analysis. **e**, Gel image of the PCR from AAV2-KKH-saCas9-sgRNA-4 edited cochlea, showing the bands of the *Mir96* locus, AAV vector DNA and miR96-ITR integration. **f**, Gel image of the PCR products, showing the bands of *Mir96* locus, AAV vectors and miR96-ITR integration of isolated hair cells from injected ears. Asterisk indicates the putative integration fragment position. **g**, MiR96-ITR integration reads from NGS of isolated hair cells from injected ears, the reference is the AAV ITR sequence. **h**, qPCR analysis of *Mir96* level in injected and uninjected cochlea (n=6). Each dot represents a independent result from 2 cochleae combined. Values and error bars reflect mean ± SD. **i**, qPCR analysis of miR96-ITR RNA level in injected and uninjected cochlea to test if there are miR96-ITR transcripts in injected cochlea (n=3). Each dot represents an independent experiment with 2 cochleae. Values and error bars reflect mean ± SD. **j**, CIRCLEseq analysis of KKH-saCas9-sgRNA-4 in *Mir96^14C>A^*^/+^ primary fibroblasts genomic DNA. **k** Quantification of indel frequency of potential off-target sites from the mouse genome. **l** Quantification of indel frequency of potential off-target sites from AAV2-KKH-saCas9-sgRNA-4 injected cochlea. Values and error bars reflect mean ± SD.

Another concern is the risk of AAV vector integration into CRISPR-induced DNA breaks^50,51^. To assess AAV vector integration, specific primers were designed for PCR and NGS analysis to detect the integration of *Mir96* and AAV ITR (Fig. 7c). In cultured HEK-293T-*Mir96^14C>A^*cells treated with different AAV dosages, AAV vector integration at the double-strand break locus became detectable at a dosage of 10^3^ vg/cell (Fig. 7d and Extended Data Fig. 6a, b). The indel frequency and the integration ratio were correlated with the increasing AAV dosage. By varying AAV dosages, we identified an optimal dosage range from 10^2^ vg/cell to 10^4^ vg/cell, where the indel frequency was near its peak while maintaining low levels of AAV vector integration in HEK cells (Fig. 7d and Extended Data Fig. 6a, b). These results suggest that by fine-tuning the AAV dosage, the potential risk of AAV vector integration can be greatly reduced while maintaining efficient genome editing.

We further performed in vivo injection and integration assay. We did not detect any integration in the whole cochlear samples (Fig. 7e), but detected minimal integration in isolated hair cells (Fig. 7f). NGS revealed that out of a total of 82,453 reads, the *Mir96* wild-type allele had 42,991 reads, while the remaining reads were associated with *Mir96^14C>A^* reads. Among the *Mir96^14C>A^* reads, indel-containing reads accounted for 25.7% (Fig. 7g). Additionally, there were 102 integration reads, making up 0.26% of the total reads (Fig. 7g). Measurement of miR96 levels using qPCR showed a decrease in miR96 levels in the injected ear (89.2± 3.0%) compared to the uninjected ear (Fig. 7h), likely due to RNA degradation after genome editing. No significant change was observed in qRNA analysis using miR96-specific and ITR-R primers (Fig. 7i), which were used to measure the miR96-ITR mRNA, indicating the absence of miR96-ITR chimera transcripts after genome editing. These results demonstrate that AAV2-mediated genome editing in the mature cochlea exhibits minimal or negligible vector integration. To further explore the possibility of eliminating AAV vector integration, a reduced dose of AAV2-KKH-saCas9-sgRNA4 (1×10^9^ vg per cochlea) was administered. After a 10-week period following the injection, no integration of ITR was detected in isolated hair cells (Extended Data Fig. 7a), as evidenced by NGS data with an average indel frequency of 12.05% (n=3) and ITR integration reads ratio below the background error (Extended Data Fig. 7b, c). These findings demonstrate that AAV vector integration rate is correlated with the administrated AAV dosage, and the dosage we used achieve the therapeutic threshold for genome editing therapy while minimize the AAV vector integration.

To assess off-target editing, CIRCLEseq was performed to identify potential off-target sites^52^. Only two off-target sites were identified from the mouse genomic DNA (Fig. 7j). Computational predictions were also used to identify potential off-target loci^53,54^. We analyzed the top ten off-target hits in mouse genome according to the cutting frequency determination (CFD) score^53^ after KKH-saCas9/sgRNA-4 editing (Extended Data Fig. 6c, d), which includes the two off-target sites identified by CIRCLEseq analysis. NGS analysis identified one off-target site with low level of indels (1.97%), while the on-target editing efficiency is up to 84.6% (Fig. 7k). The off-target editing was in the intergenic region (Gm19782-Fam135b), which means unlikely to disrupt any gene function. Using the isolated hair cells from injected mice, we did not detect any off-target indels (Fig. 7l). Together, these results show that the delivery of AAV2-KKH-saCas9-sgRNA-4 into *Mir96^14C>A/+^* cells results in minimal off-target modification, and the hearing-related phenotypes are the result from on-target editing.

### A dual AAV2 system enables targeting multiple human *MIR96* mutations

As the *Mir96* seed sequence is 100% conserved across mammalian species, the gRNA designs that target mouse mutations can be directly used to target orthologous human mutations. In addition to the +14C>A mutation, there are two known dominant mutations in the *MIR96* seed region, *MIR96* 13G>A and *MIR96* 15A>T, in humans^11,17^. We designed gRNAs for each of the two mutations (Fig. 8a). Cas9 and multiple sgRNA cassettes are too big to fit in single AAV, we then created a dual AAV set consisting of AAV-U1A-spCas9-polyA with three nuclear localization signals (NLSs) and sgRNA cassettes to target all three mutations (14C>A, 13G>A and 15A>T), AAV-sgmiR96-master (Fig. 8b, c). To validate the dual AAV set, human cell lines containing the three *MIR96* mutations were generated using the PiggyBac system, and each cell line was transfected with AAV-U1A-spCas9-polyA/AAV-sgmiR96-master. NGS revealed robust indel formation in all three *MIR96* mutation lines, with a similar editing efficiency (Fig. 8d-g and Extended Data Fig. 8a-f). Less than 1% of the reads had indels detected at the *MIR96* locus (Fig. 8e). The data support specific disruption of each mutant allele by its corresponding gRNA. We also analyzed top 10 potential off-target sites in human genome and NGS showed no indels after Cas9/sgRNA transfection (Fig. 8h, j and Extended Data Fig. 8g). These findings suggest that the combination of AAV-U1A-spCas9-polyA/AAV-sgmiR96-master can effectively target each of the three known human *MIR96* mutations and holds promise for treating dominant hearing loss caused by *MIR96*-related mutations. The design of the AAV-sgmiR96-master system expands the targeting scope and simplifies the efficacy and safety evaluation of the AAV delivery system.

**Fig. 8.**
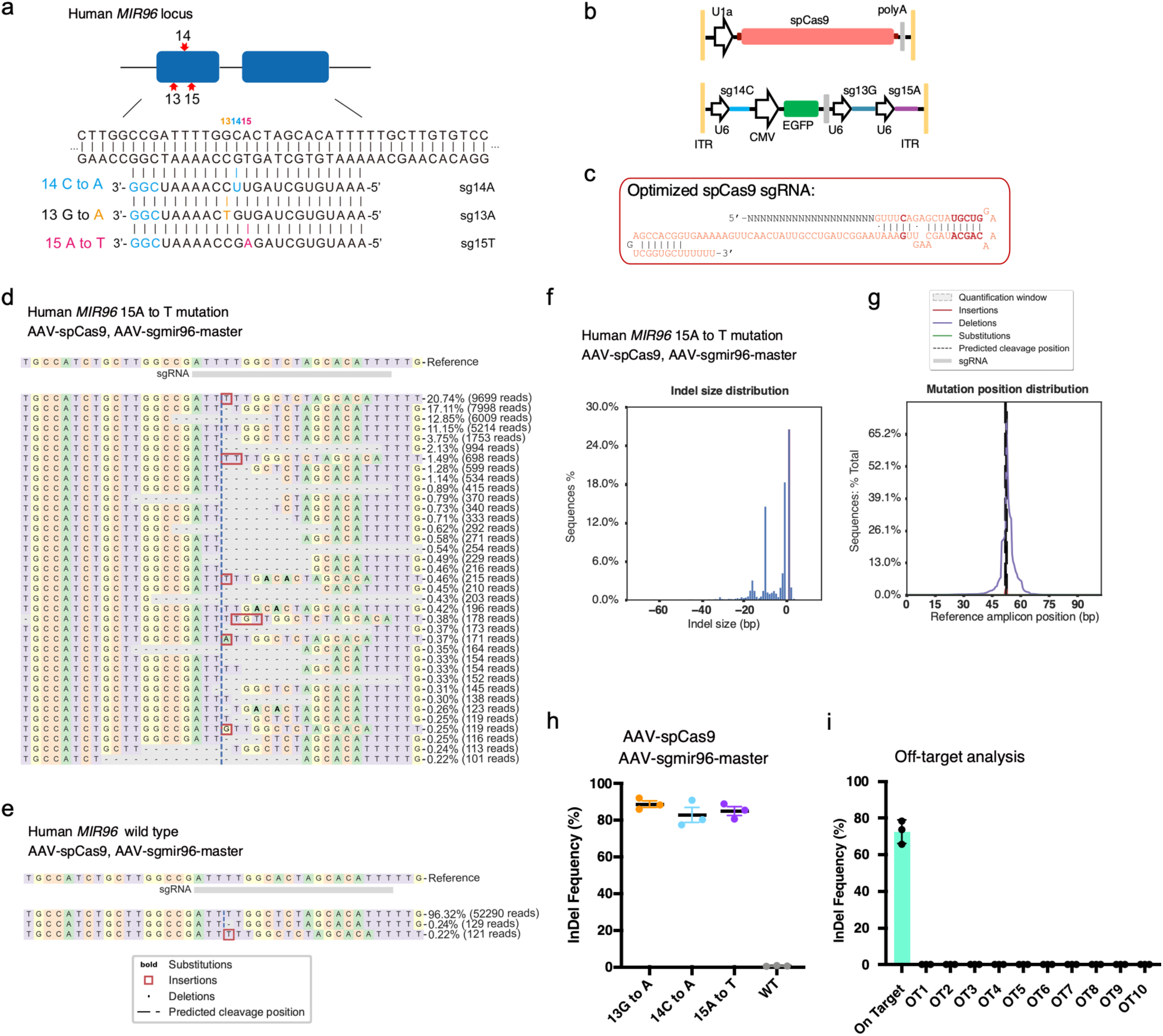
A dual AAV carrying multiple gRNAs targeting all known *Mir96* seed region mutations in human cells specifically and efficiently. **a**, Sequence information of the human *MIR96* locus and the sgRNAs design to target three known seed mutations. Red: mutatio site. Blue: the protospacer adjacent motifs (PAMs) nucleotides. **b**, Schematic view of dual AAV constructions. One contains “U1a-spCas9-polyA” cassette, the other contains three U6-sgRNA cassettes. **c**, Sequence of the optimized spCas9 sgRNA, the bold lettersindicate the changes compared to the unmodified sequence. **d, e**, Representative NGS results from spCas9/sgmiR96-master edited HEK-miR96 (15A to T) cells and spCas9/sgmiR96-master edited HEK293T wild type cells. **f, g**, Indel profiles from spCas9/sgmiR96-master edited HEK-miR96 (15A to T) cells. Minus numbers represent deletions, plus numbers represent insertions. **h**, The indel frequency in the HEK-miR96 mutation cell lines and wild-type HEK293T cells after genome editing using spCas9/sgmiR96-master. Each dot represents an independent experiment. Values and error bars reflect mean ± SD. **i**, Off-target analysis in spCas9/sgmiR96-master edited HEK-miR96 (15A to T) cells. Genomic DNA was pooled from three independent biological replicates for sequencing. Values and error bars reflect mean ± SD.

## DISCUSSION

With the goal of developing a clinical treatment for human genetic hearing loss of DFNA50, we conducted genome editing in adult *Mir96^14C>A/+^* cochleae and successfully improved auditory function in the long term. miRNAs are involved in multiple physiological and pathological inner ear processes^55^, and have been linked to different types of hearing loss, including deafness related to hair cell development ^8,56^, age-related hearing loss^57^, noise-induced hearing loss^58^, and inner ear inflammation^59^. In humans, heterozygous *MIR96* mutations result in late onset non-syndromic progressive hearing loss, indicating the potential loss of sensory hair cell identity and subsequent dysfunction in the mature cochlea, thereby offering an opportunity for genetic intervention in adult patients.

Specific and efficient sgRNAs and genome editors, spCas9-sgRNA-1 and KKH-saCas9-sgRNA-4 were identified and optimized for targeting the *Mir96^14C>A^* mutation. AAV2 was chosen as the delivery vehicle for its efficient transduction in inner and outer hair cells. Genetic disruption of the *Mir96^14C>A^* allele leads to improved survival of hair cells and the preservation of auditory function in *Mir96^14C>A/+^* mice. We conducted injections of AAV2 into adult mice across various age groups. The therapy was effective in improving long-term hearing preservation in both pre-symptomatic and symptomatic stages of hearing loss in *Mir96^14C>A/+^* mice. Notably, we observed transient expression of KKH-saCas9 in the inner ear, as the CMV promoter became silenced over time in vivo. This temporary expression provides a controlled and regulated editing process, which enhances the safety and precision of the therapy. Additionally, we demonstrated that fine-tuning the dosage of AAV helps minimize the risk of AAV-mediated integration and improves the safety profile of the therapy. These findings greatly enhance the safety profile of in vivo genome editing therapy. The study also introduces a dual-AAV system capable of targeting all known human *MIR96* seed region mutations, making it a promising candidate for treating various *MIR96* dominant mutation-caused genetic hearing loss conditions. Our study highlights the editing efficiency, safety, long-term therapeutic efficacy of genome editing in treating hearing loss due to *Mir96* mutations in adult animal models and provides a promising path for clinical applications.

The selection of the most suitable editor and delivery vehicle is crucial in developing a genome editing therapy for specific genetic disorders. In this study, we chose Cas9 nucleases to disrupt the mutated allele of *Mir96^14C>A/+^* due to its gain-of-function inheritance pattern^18^ meaning that the resulting indels would disable the expression of the disease-causing mutant *Mir96*. While prime editing has the potential to correct the *Mir96* mutation, our in vitro data showed modest efficiency at the *Mir96* locus, requiring further optimization. AAV2 was selected as the preferred delivery vehicle for CRISPR due to due to its specific targeting of inner and outer hair cells in the cochlea of adult animals^39^. Besides, AAV2 serotype remains the most used throughout the study period and has the most safety and efficacy evidence, with the most completed clinical trials^60^.

One of the major concerns associated with AAV-mediated CRISPR genome editing in vivo is the potential for off-target editing caused by prolonged expression of Cas9/sgRNA^61^. High-fidelity versions of CRISPR systems^35,62,63^ can be applied to improve specificity. For example, we have demonstrated LZ3 Cas9 can target the *Mir96^14C>A^* allele robustly and specifically. In addition, minimizing genome editing in unintended tissues and cell types is also an important way to enhance the safety of CRISPR-based therapeutics. AAV2 can specifically deliver Cas9/sgRNA into hair cells through local injection, reducing the likelihood of editing unwanted cells or tissues. Besides, the use of the CMV promoter for KKH-saCas9 expression helps minimize off-target editing as it tends to become silenced over time in vivo^47–49,64–66^. We demonstrated that CMV-driven KKH-Cas9 was only transiently expressed and silenced in 3 months in the cochlea. Another concern is the risk of vector integration at DNA double-stranded breaks (DSBs)^50,51^. The integration ratio of AAV varies across different genomic loci, tissues, and cell types^50^. By focusing on hair cell-limited genome editing, we were able to reduce the occurrence of vector integration, as demonstrated in our results. Furthermore, we found that a higher dose of AAV increased the chance of vector integration, underscoring the importance of carefully fine-tuning the dosage. To advance clinical studies, it will be necessary to dissect the AAV injection dosage to identify the threshold for achieving robust genome editing while minimizing AAV vector integration.

MicroRNA-96 serves as a key regulator and plays a crucial role in hair cell development and hearing maintenance. Our study demonstrated in vivo genome editing in adult *Mir96^14C>A/+^* mice, in both pre-symptomatic and symptomatic stages of hearing loss, can improve auditory function and prevent long-term deterioration. Moreover, our study highlights the importance of genetic intervention in adult mice of different age groups to determine the optimal treatment window, which has implications for future clinical studies. Moving forward, it will be necessary to conduct further investigations in large animals such as non-human primates to evaluate the efficacy of targeted genome editing, determine the ideal AAV administration dosage, and identify the most effective intervention time window for maximum therapeutic benefit. Our work holds significant potential in advancing the clinical application of genome editing therapy for treating genetic inherited hearing loss.

## Methods

### Animals and surgery

The *Mir96^14C>A^* allele (official symbol *Mir96^tm^*^3^*^.1Wtsi^*, referred to here as *Mir96^14C>A^*) was generated at the Wellcome Sanger Institute as part of the Mouse Genetics Project, by targeting of the BEPD0003_D07 ES cell of C57BL/6N origin and was maintained on the same genetic background. The selection cassette was removed by FLP recombinase, leaving a single downstream FRT site. All procedures for generating the mutant mouse were carried out in accordance with UK Home Office regulations and the UK Animals (Scientific Procedures) Act of 1986 (ASPA) under UK Home Office licences, and the study was approved by the Wellcome Sanger Institute Ethical Review Committee. After shipping, all animals were bred and housed in Mass Eye and Ear Infirmary. All studies involving animals were approved by the HMS Standing Committee on Animals and the Mass Eye and Ear Infirmary Animal Care and Use Committee. All mice were housed in a room maintained on a 12h light and dark cycle with ad libitum access to standard rodent diet.

*Mir96^14C>A/+^* and wild type (WT) mice of either sex were anesthetized using intraperitoneal (i.p.) injection of ketamine (100mg/kg) and xylazine (10mg/kg). The post-auricular incision was exposed by shaving and disinfected using 10% povidone iodine. The AAV2-CMV-KKH-saCas9-sgRNA4 of two titers was injected into the inner ears of *Mir96^14C>A/+^* mice. The AAV2-GFP was injected into the inner ears of WT mice. The total volume for each injection was 1∼1.2 μl virus per cochlea.

### Plasmid construction

pMax-spCas9 is obtained from previous article^67^, U6-sgRNA sequence was also obtained from PX458 (Addgene 48138)^68^, other Cas9 nuclease, including scCas9++ (Addgene 155011)^32^, KKH-saCas9 (Addgene 70707)^33^, sauriCas9 (Addgene 135964)^34^, LZ3 Cas9 (Addgene 140561)^35^ cDNA are acquired from addgene. Vectors for in vitro screening were constructed via Gibson assembly (NEB, E2611S). KKH-saCAs9 AAV vectors are based on PX601 (Addgene 61591)^69^. AAV-U1a-opti-spCas9-PA (Supplementary Fig. 2) is based on Addgene 121507^70^. AAV-sgmir96-Master (Supplementary Fig. 3) is based on pAAV-U6-sgRNA-CMV-GFP (Addgene 85451)^71^. Plasmids encoding recombinant AAV (rAAV) genomes were cloned by Gibson assembly. All plasmids were purified using Plasmid Plus Miniprep or Maxiprep kits (Qiagen).

### AAV production

AAV vectors were produced by Mass eye and ear infirmary vector core (Boston, MA, USA). AAV plasmids containing “CMV-KKH-SaCas9-PA” cassette, “U6-sgRNA-4” or “U6-sgCtrl” cassette was sequenced before packaging (MGH DNA Core, complete plasmid sequencing) into AAV2/2 (Supplementary Fig. 1). Vector titer was 5.76 × 10^12^ vg/ml for KKH-saCas9/sgRNA-4 and 4.55× 10^12^ vg/ml for KKH-saCas9/sgCtrl as determined by qPCR specific for the inverted terminal repeat of the virus.

### Isolation and culture of Primary Fibroblasts from Mice

*Mir96^14C>A/+^,* Ai14 (CAG-STOP-tdTomato) and wild type mice were euthanized and cleaned with 70% ethanol. The dorsal skin (∼1 cm diameter) of the mice was collected and rinsed with DPBS, subcutaneous fat was removed by forceps. Subsequently, the samples were cut into small fragments and incubated with Dispase II (Sigma-Aldrich, USA) for overnight at 4 °C. The dermal layers of the skin were separated using forceps and further subjected to incubation with type I collagenase (1 mg/ml Gibco, USA) for 2 hours at 37 degrees. The resulting cell suspension was strained using a 40-micron strainer and centrifuged at 950 rpm to obtain the cell pellet. The cell pellet was seeded in T-75 flask containing DMEM high glucose media (Gibco, USA) containing 10 % FBS (Gibco, USA). Fibroblasts were cultured for about 2–3 days to reach ∼90% confluence, then passaged in T75 flasks with TrypLE Express and cultured in DMEM: F12 medium (ThermoFisher) with 10% fetal bovine serum (FBS) supplemented with GlutaMax (ThermoFisher).

### Construction of *Mir96^14C>A^* cell line using PiggyBac

Mouse *Mir96^14C>A^* (∼0.6 kb) harboring the +14C>A mutation was amplified by PCR from *Mir96^14C>A^*^/14C>A^ mouse genomic DNA. The PCR products were cloned into the PiggyBac donor backbone (PB-CAG-mNeonGreen-P2A-BSD-polyA) using Gibson Assembly. The constructed donor plasmid was co-transfected with PiggyBac transposon vector (PB210PA, System Biosciences) into HEI-OC1 cells. Cells were cultured and selected in the medium containing 10 µg/mL Blasticidin for 2 weeks. For human *MIR96* +14C>A fragment, mutation was introduced by PCR and cloned into the same PiggyBac donor backbone using Gibson Assembly. Cells were transfected by PiggyBac plasmids and selected by Blasticidin for 2 weeks. Successful insertion was confirmed by PCR and sequencing analysis. Clones from the positive *Mir96^14C>A^* selection were expanded for subsequent studies.

### Genome editing in vitro

We performed nucleofection using LONZA 4D-Nucleofector. Cells were digested by Trypsin-EDTA (0.05%) (Thermo Fisher) and future dispersed into single cells. 100,000 cells were resuspended in 20μl P3 reagent of the P3 Primary Cell 4D-Nucleofector® X Kit S (Lonza V4XP-3032). 1 μg total plasmid was used for a single nucleofection event and nucleofected by program EH-100. Cells were not sorted. 5 days after the nucleofection, cells were lysed by QuickExtract™ DNA Extraction Solution (Lucigen) to extract genomic DNA. Genomic PCR was carried out using NEBNext® Ultra™ II Q5® Master Mix (NEB, M0544S) to amply *Mir96* loci, 400∼800 ng of purified PCR product were used for next generation sequencing (NGS), samples were sequenced and analyzed to detect CRISPR variants from NGS reads using CRISPResso2 (http://crispresso.pinellolab.org/submission) as instructed^72^. The sgRNA protospacer sequences can be found in Supplementary Table 3.

### Hair cells isolation and NGS analysis

Cochleae were harvested with the sensory epithelia dissociated using needles under the microscope (Axiovert 200M, Carl Zeiss). Inner ear tissue was immersed in 1 μM FM 1-43FX (ThermoFisher, F35355) dissolved in DPBS (ThermoFisher) for 15s at room temperature in the dark, then washed by DPBS. The sensory epithelia were treated with 100μl 0.05% trypsin-EDTA (ThermoFisher, 25300054) for 10-20 min. During incubation, the tissue was carefully dispersed into small cell clusters or single cells using a 200μl Eppendorf pipette tip. Cells were then transferred into 6-well plate and placed under a fluorescent microscope (ZEISS) equipped with a camera. Cells with FM 1-43FX dye were collected by a 10μl Eppendorf pipette tip. Above 300 cells were collected from single cochlea. Hair cells were transferred into 200μl PCR tube, and centrifuged for 5 min, 200 rcf. Supernatant were carefully discarded and the isolated hair cells were lysed by 5μl QuickExtract™ DNA Extraction Solution (Lucigen) and incubated in 65°C for 6 min then 98°C 3 min. All 5μl of the cell lysis were used for each Genomic PCR amplification using NEBNext® Ultra™ II Q5® Master Mix (NEB, M0544S). PCR program is 1 cycle: 98°C 5min; 42 cycles: 98°C 15s, 60°C 20s, 72°C 10s; 1 cycle: 72°C 4min; 4°C. PCR products visualization, purification and NGS analysis are the same with that described above.

### Hearing Function Testing

*Mir96^14C>A/+^* mice of either sex were anesthetized using intraperitoneal (i.p.) injection of ketamine (100mg/kg) and xylazine (10mg/kg). For ABR measurements, subcutaneous needle electrodes were inserted at the vertex, ventral edge of the pinna (active electrode), and a ground reference near the tail. The mice were placed in a sound-proof chamber and exposed to 5-ms tone pips delivered at a rate of 35/s. The response was amplified 10,000-fold, filtered with a band-pass of 100 Hz to 3 kHz, digitized, and averaged using 1,024 responses at each sound pressure level (SPL). The sound level was elevated in 5 dB steps from 20dB up to 90dB SPL, with stimuli ranging from 5.66-45.24 kHz frequencies (in half-octave steps). The “threshold” and wave 1 amplitude were identified as described previously^19,20^. During the same recording session, DPOAEs were measured under the same conditions as for ABRs. Briefly, two primary tones (f2/f1=1.2) were set with f2 varied between 5.66 and 45.24 kHz in half-octave steps. Primaries were swept from 20dB SPL to 80dB SPL (for f2) in 5-dB steps. Thresholds required to produce a DPOAE at 5dB SPL were computed by interpolation as f2 level.

### Immunofluorescence staining

Cochleae, both injected and non-injected, were harvested following CO_2_ inhalation as a means of animal euthanasia. Temporal bones were fixed in 4% paraformaldehyde at 4°C overnight and subsequently decalcified in 120 mM EDTA for a week. Then the organ of Corti was dissected for whole-mount immunofluorescence. The dissected tissues were blocked with a blocking solution (PBS with 8% donkey serum and 0.3-1% Triton X-100) for 1 hour at room temperature. Subsequently, the specimens were subjected to overnight incubation with the primary antibodies: anti-MYO7A (#25-6790, Proteus BioSciences), anti-GFP (ab13970, Abcam). After three rinses with PBS, the tissues were incubated with the corresponding secondary antibodies for 1 hour. Finally, all specimens were mounted with VECTASHIELD antifade mounting medium containing DAPI (VECTOR LABORATORIES, #H-1200). Images were taken with a Leica SP8 confocal laser scanning microscope (Leica Microsystems, Germany). For hair cell counting, MYO7A-positive hair cells per 100μm length were calculated in the apex, middle, and middle-base turns of cochleae. We counted from three independent cochleae.

### Scanning Electron Microscopy

Following cochlea dissection, the harvested tissues were placed in 2.5% glutaraldehyde solution in 0.1 M cacodylate buffer (EMS) supplemented with 2 mM CaCl2. The immersion was performed for 1.5-2 hours at room temperature on a tissue rotator. Subsequently, the samples were rinsed three times with distilled water. The samples underwent three 10-minute rinses with 0.1 M sodium cacodylate buffer. Next, they were treated with 1% osmium tetroxide for a duration of one hour, followed by another three 10-minute rinses with distilled water. The samples were then subjected to treatment with saturated thiocarbohydrazide in distilled water for 30 minutes. This cycle of treatments (one-two-one-two-one) was repeated.

For dehydration, the samples were transferred to 20 ml scintillation vials containing 2 ml of distilled water. The vials were supplemented with 50 µL of 100% ethanol, with the volume doubled every 10 minutes until the vial was full. At that point, the samples were transferred to 100% ethanol. Subsequently, the samples were dried using liquid CO2 in a Tousimis Autosamdri 815 to reach the critical point. Finally, the samples were mounted onto aluminum specimen stubs using carbon tape, spatter-coated with a 4.5 nm layer of platinum using a Leica EM ACE600 and analyzed using a Hitachi S-4700 scanning electron microscope (SEM).

### RNA isolation and qRT-PCR

Total RNA was extracted from inner ear tissue using the ReliaPrep RNA Tissue Miniprep System (Promega, z6111). Then first-strand cDNA was produced using ProtoScript® II First Strand cDNA Synthesis Kit (NEB, E6560s) with random primers, following the manufacturer’s instructions. STEM-LOOP qRT-PCR were used to measure *Mir96* level, first-strand cDNA was produced using *Mir96* specific primers (rtmri96: CTCAACTGGTGTCGTGGAGTCGGCAATTCAGTTGAGCAAAAATGTG). Real time quantitative PCR was performed using Power SYBR Green PCR Master Mix (Applied Biosystems, 4368708) on the ABI QuantStudio 3 Flex Real-Time PCR System (Applied Biosystems). qmiR96-F: TCGGCAGGTTTGGAACTAGCAC; qmiR96-R: CTCAACTGGTGTCGTGGA.

### Western blotting

After dissection of cochleae, inner ear tissue samples were lysed using by RIPA Lysis and Extraction buffer (ThermoFisher, 89900) containing protease inhibitor (ThermoFisher) for 30 min on ice. Protein was quantified and 80 μg of each lysate were loaded per lane of a NuPAGE™ 4-12% Bis-Tris Protein Gel (Thermo Fisher Scientific, NP0335PK2). Samples were separated on 200V for 35 min in the mini gel tank (ThermoFisher). Protein were then transferred to Nitrocellulose Blotting Membranes (PALL, P66485) at 200 mA for 2 hours. The membranes were blocked in 5% evaporated milk in Tris-based saline with Tween 20 (0.05% TBST) for 1 hour and followed by incubating with primary antibodies overnight at 4 °C (Anti-HA: Cell Signaling Technology, 3724S; anti-GAPDH: Thermo, MA5-15738). After three rinses with TBST, NC membranes were incubated with HRP conjugated secondary antibodies (Thermo Fisher Scientific) for 2 hours at RT. Protein bands were visualized using SuperSignal West Pico PLUS Chemiluminescent Substrate (ThermoFisher, 34577) and blot image were captured by ChemiDoc Imaging system (BioRad). Uncropped images can be found in Supplementary Fig. 4.

### AAV vector integration assay

HEK 293T-*Mir96^14C>A^* cells were treated with AAV2-CMV-KKH-saCas9-sgRNA4 of different dosages from 1 to 10^7^ genomic copies per cell. Control cells were transduced with AAV2-CMV-KKH-saCas9-sgCtrl, 10^5^ genomic copies per cell. Cells were collected 7 days later and genomic DNA was isolated. Primers P1-F/ITR-R were used for detecting of AAV vector integration. For in vivo integration detection, Primers P1-F/P2-R and primers P1-F/ITR-R were used to amply genomic fragment from isolated hair cells, then PCR products were merged together for NGS analysis. Primers premir96-F/ITR-R were used in the qRT-PCR study to detect mir96-ITR transcription. All primers used for AAV vector integration assay were listed in Supplementary Table. 1.

### Off-target analysis

To identify off-target sites, the CIRCLE-seq was performed essentially as previously described with the minor modifications. Briefly, 50 μg genomic DNA were purified from HEI-OC1 cells. Genomic DNA was sheared (NEB Ultra II FS kit, E7805) and circularized. In vitro cleavage reaction of circularized genomic DNA is performed with KKH-saCas9/sgRNA-4. Then the sequencing libraries were prepared and sequenced on an Illumina Miseq as previously described^52^. Off-target sites were identified with the standard pipeline. The negative control sample were treated with the Cas9 alone to assess background.

Besides, we listed top10 potential off-target sites according to the cutting frequency determination (CFD) score^53^. We designed PCR primers on the two flanks of each sgRNA target sequences for amplifying 250-280 bp DNA fragments. Then amplified fragments were purified for NGS to identify whether there is any off-target mutation. All primers used for off-target analysis were listed in Supplementary Table. 2.

### Statistical analysis

The number of biological and technical replicates, and parameters are indicated in corresponding Figure legends. Data acquisition were performed by investigators blinded to experimental groups. Statistical analysis were performed using GraphPad Prism 8. The student’s t test was used to calculate statistical significance between two groups. The p values < 0.05 were considered as significantly different. **** p < 0.0001, *** p < 0.001, ** p < 0.01, * p < 0.05.

### Reporting summary

Further information on research design is available in the Nature Research Reporting Summary linked to this article.

## Data availability

The data supporting the results in this study are available within the paper and its Supplementary Information. AAV genome sequences are provided in the Supplemental Data. All data are available within this study are available from the corresponding author upon reasonable request. The mutant mice are available from the European Mutant Mouse Archive (EMMA).

## Funding

This work was supported by U.S. NIH R01 DC016875, R01 DC019404, UG3TR002636, U24HG010423, UH3TR002636, Curing Kids Massachusetts Eye & Ear and Ines-Fredrick Yeatts Fund (to Z.-Y.C.), NIH R01 DC019404 (to X.L. and Z.-Y.C.), R01DC012115, R01DC005575, and DOD RH220053 (to X.L.). This work was funded in part by the Wellcome Trust (098051, 100669, 089622; KPS). For the purpose of Open Access, the author has applied a CC BY public copyright license to any Author Accepted Manuscript (AAM) version arising from this submission.

We thank the Wellcome Sanger Institute Mouse Genetics Project for generating and providing the *Mir96* 14C>A mutant mouse.

We thank the Harvard Medical School Electron Microscopy Facility for their help on the Scanning Electron Microscopy Imaging.

## Author contributions

Conceptualization: ZYC, WZ, WD

Methodology: WZ, WD

Experiment: WZ, WD, APR, AA, SS, YK, WW

Data analysis: all authors Supervision: ZYC

Manuscript writing: WZ, WD, ZYC

Manuscript review and editing: WZ, WD, APR, AA, SS, YK, WW, YS, XL, MAL, KPS, ZYC

## Extended Data

**Extended Data Fig. 1.**
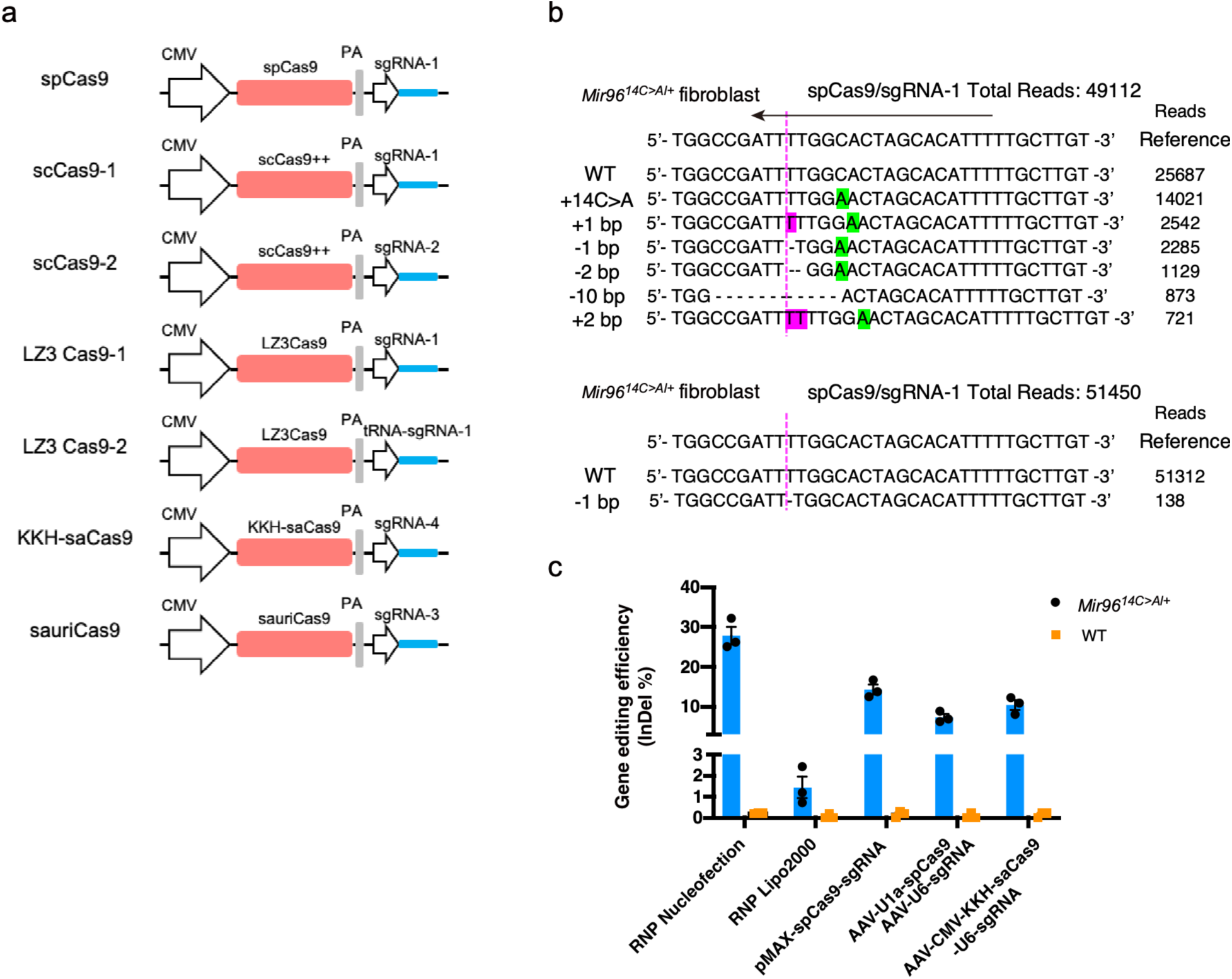
Targeting *Mir96^14C>A^* mutation with different CRISPR nuclease systems. **a**, Schematic diagram of plasmid constructions for different CRISPR systems used for in vitro screening. **b**, Representative reads of the NGS from spCas9/sgRNA-1 edited *Mir96^14C>A/+^*and wild-type primary fibroblasts. Magenta Dotted Lines indicate the double stranded DNA cutting site. Green indicates the mutant nucleotide. **c**, The indel frequency in *Mir96^14C>A/+^* and wild-type primary fibroblasts after editing by different delivery methods and editors. Each dot represents an independent experiment. Values and error bars reflect mean ± SD.

**Extended Data Fig. 2.**
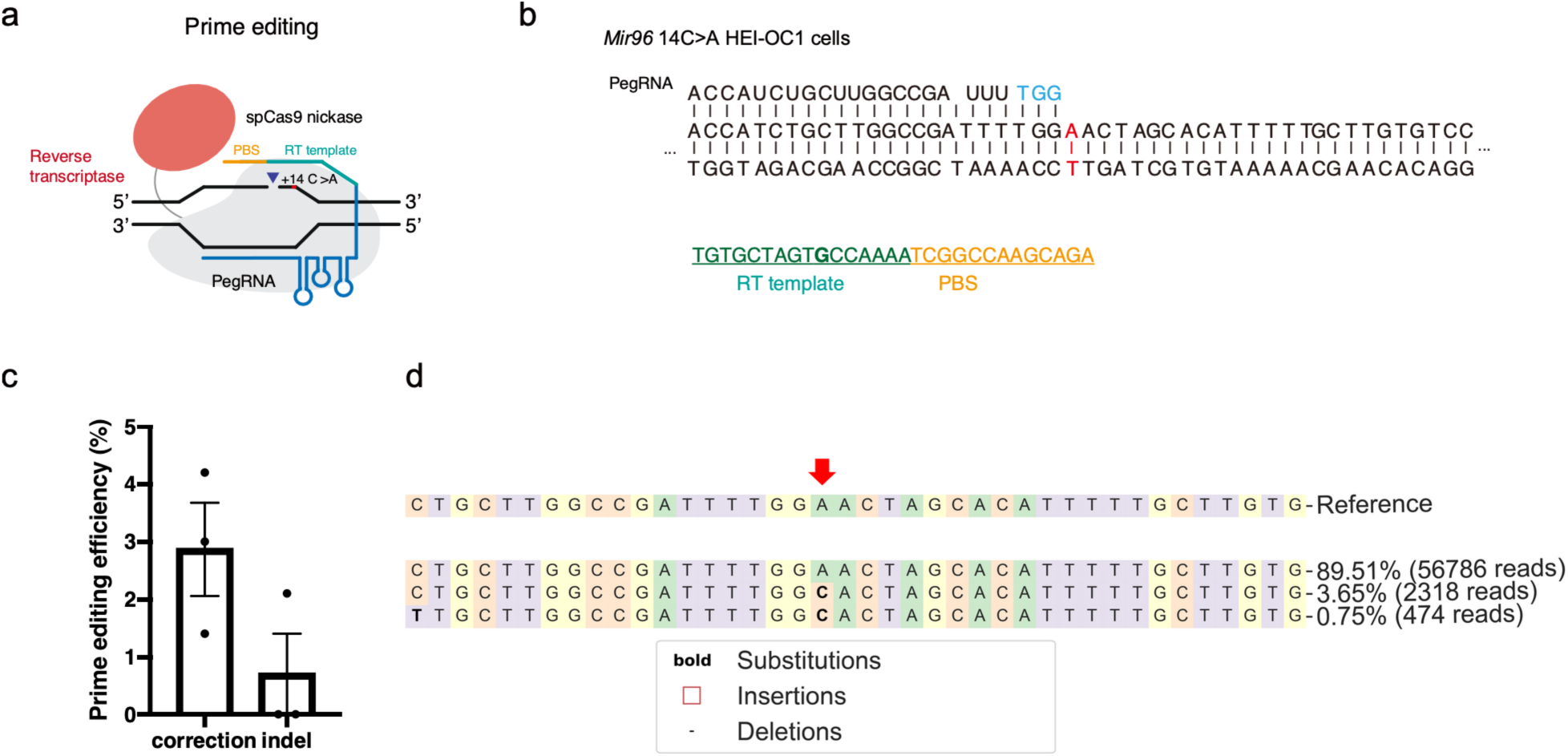
The correction of *Mir96^14C>A^* mutation using prime editing in human cells. **a**, Schematic overview of the prime editing system. **b**, Sequence of the *Mir96* +14 C>A mutation locus and the prime editing design. The mutation nucleotide in the *Mir96^14C>A^* allele is displayed in red. The protospacer adjacent motifs (PAMs) nucleotides of the pegRNA are displayed in blue. **c**, A to C correction frequency after prime editing. Each dot represents an independent experiment. Values and error bars reflect mean ± SD. **d**, Representative NGS result after prime editing, showing the corrected reads. Red arrow indicates the *Mir96* +14 C>A mutation nucleotide.

**Extended Data Fig. 3.**
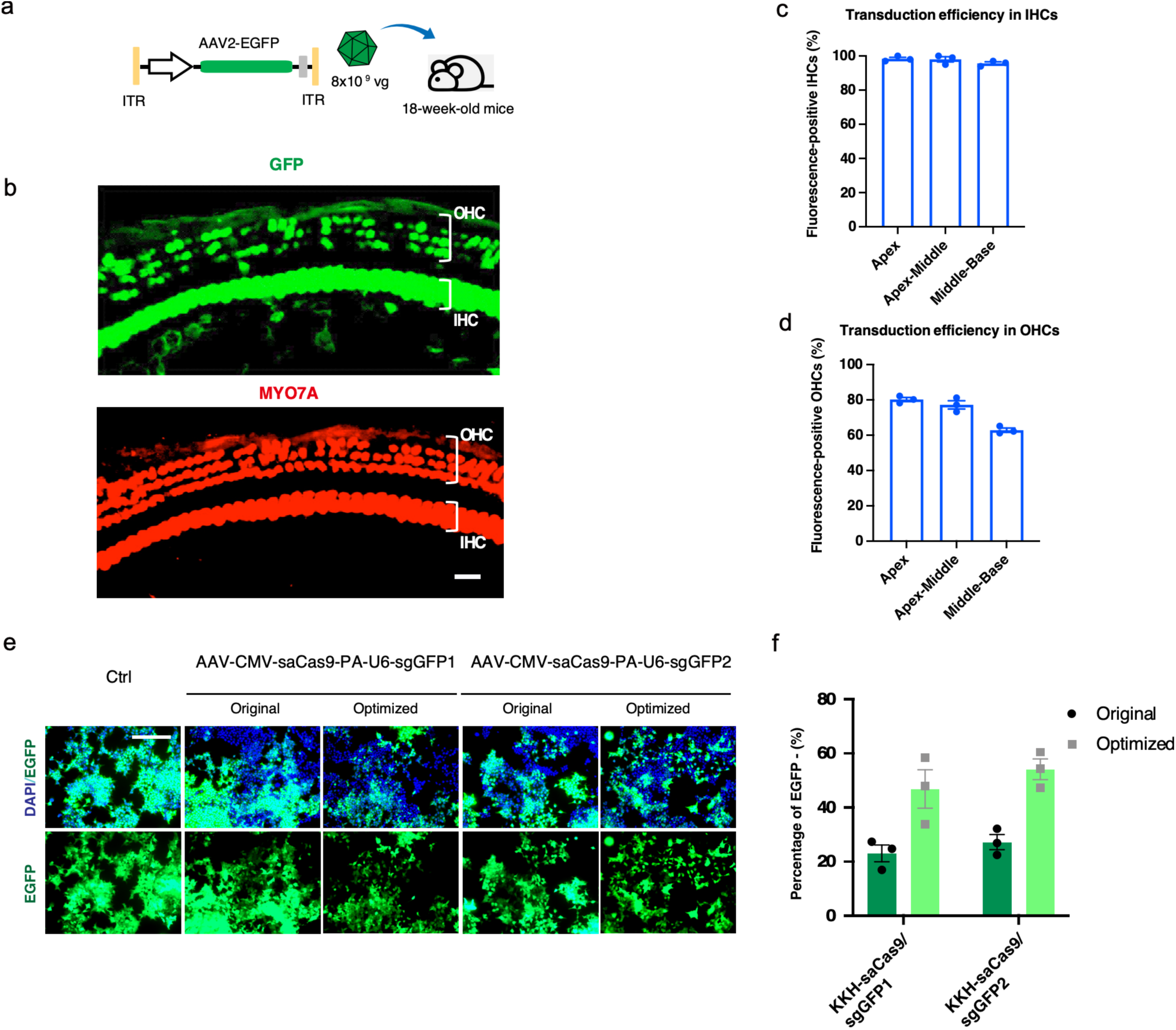
Genome editing efficiency of optimized AAV vectors. **a**, Schematic diagram of AAV2 constructions for GFP delivery used for in vivo delivery into adult mouse cochleae. **b**, Representative confocal images of AAV2 transduction in the apical region of adult cochlea. Hair cells were labeled with MYO7A (red) and AAV was labeled with GFP (green). Scale bar, 20μm. **c,d,** Transduction efficiency of IHCs (c) and OHC (d) by AAV2-GFP across different turns of the cochlea (Apex, Apex-Middle, and Middle-Base). Time of imaging: 22 weeks of age. Values and error bars reflect mean ± SD, n=3. Each dot represents an independent experiment. **e**, Representative fluorescence images of HEK-GFP cells after gemome editing of unmodified and optimized KKH-saCas9/sgRNA targeting GFP. GFP negative cells are the edited cells. **f**, Editing efficiency shown by the percentage of GFP negative cells after genome editing. Each dot represents an independent experiment. Values and error bars reflect mean ± SD.

**Extended Data Fig. 4.**
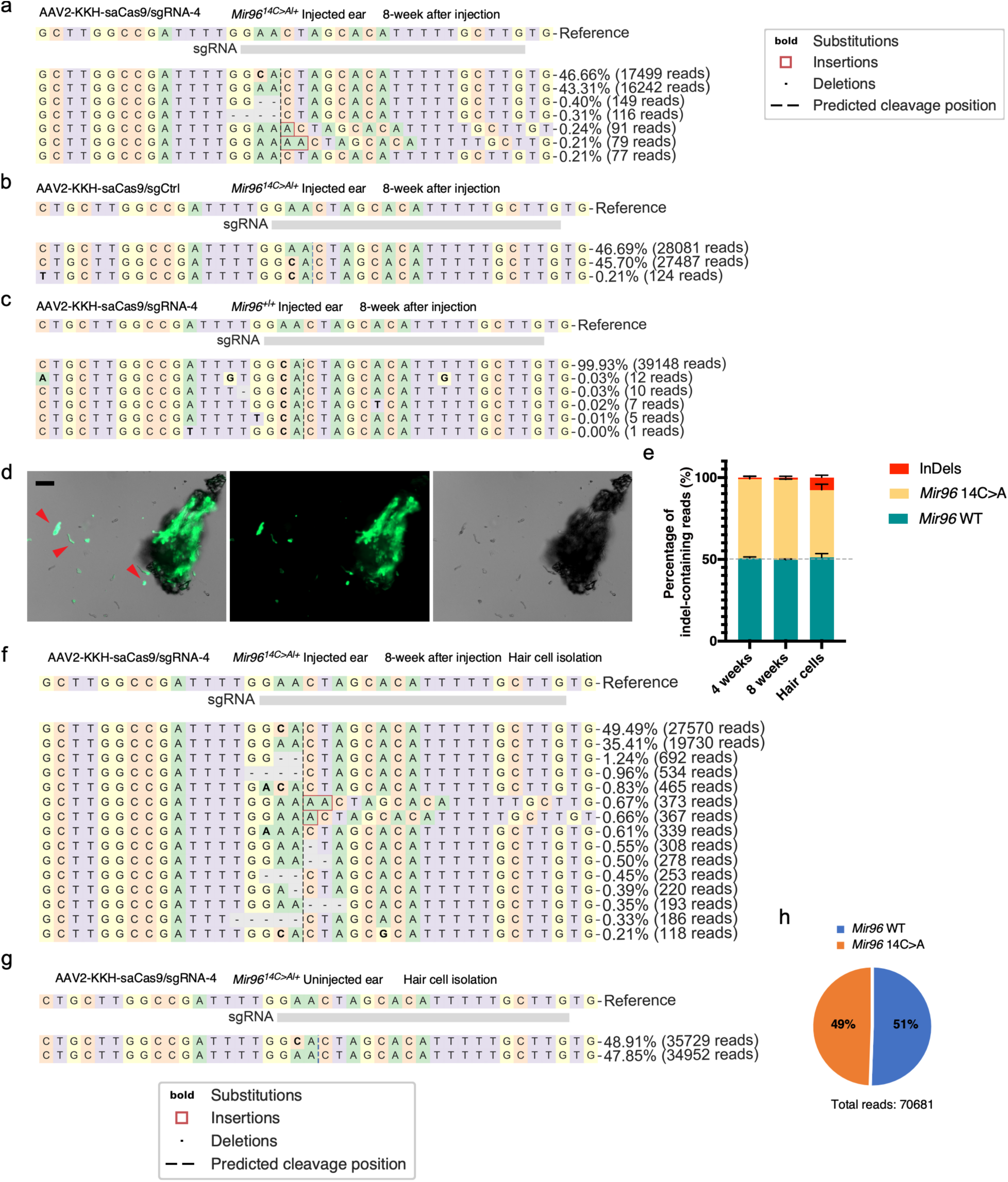
Genome editing at *Mir96^14C>A^* locus in adult *Mir96^14C>A/+^* mice. **a-c**, Representative NGS results of AAV2-KKH-saCas9-sgRNA-4 injected ears from *Mir96^14C>A**/+**^* mice and *Mir96*^+/+^ mice, and AAV2-KKH-saCas9-sgCtrl injected ears from *Mir96^14C>A**/+**^* mice. **d**, Representative images of FM1-43FX labeled hair cells after digestion of cochlear tissues. FM1-43FX labeled hair cells are shown in green. Arrowheads point to the GFP^+^ hair cells that were picked for NGS. **e**, Quantification of the percentage of indel-containing reads in the NGS results of AAV2-KKH-saCas9-sgRNA-4 edited cochlea samples and isolated hair cells lysis. Values and error bars reflect mean ± SD. **f, g**, Representative NGS results of the isolated hair cells from AAV2-KKH-saCas9-sgRNA-4 injected and uninjected ears of *Mir96^14C>A**/+**^* mice.

**Extended Data Fig. 5.**
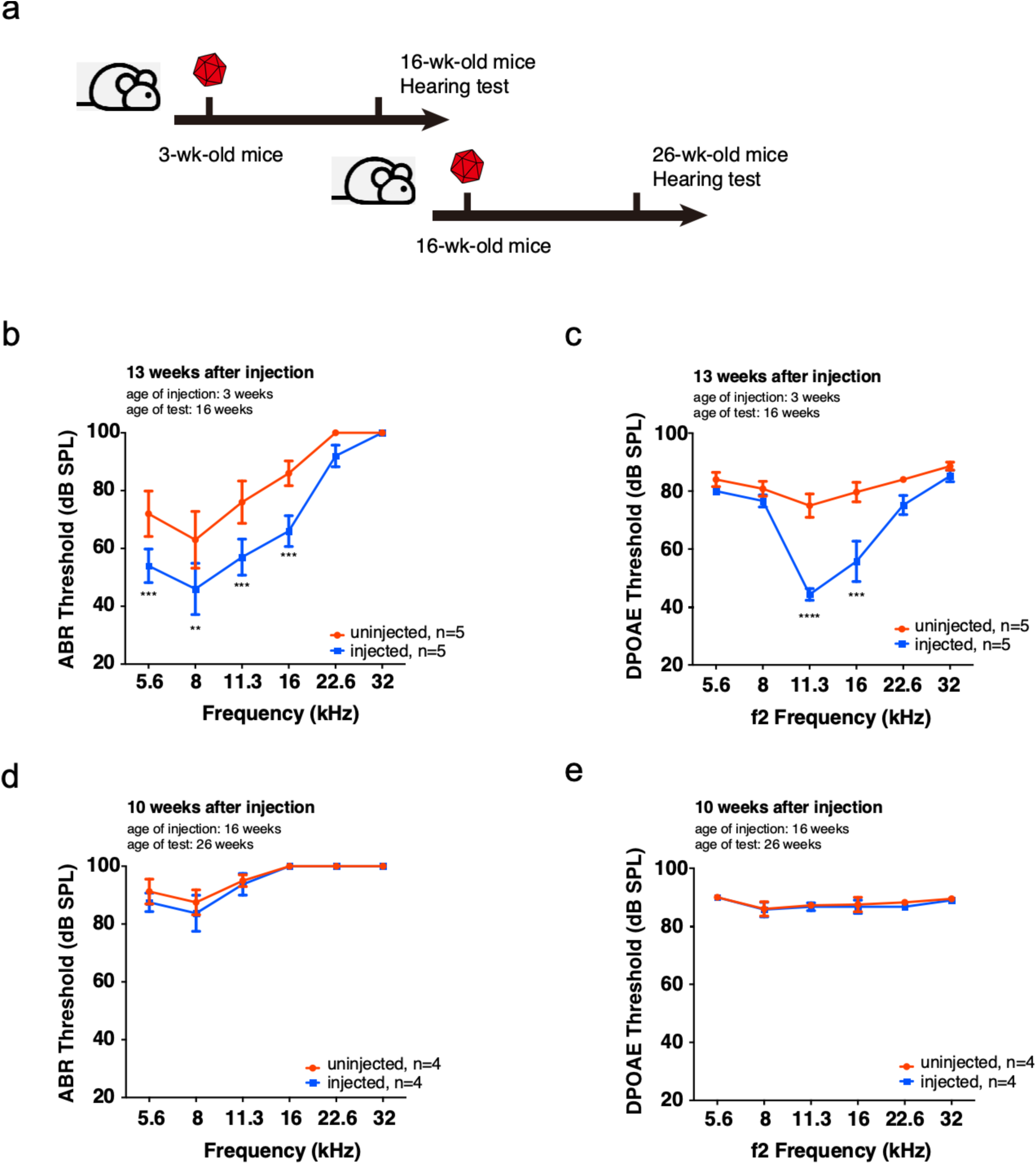
Age-dependent treatment outcome by editing in *Mir96^14C>A/+^* mice. **a,** The experimental design of inner ear injections of AAV2-CMV-KKH-saCas9-sgRNA-4 in 3-week-old and 6-week-old *Mir96^14C>A**/+**^* mice followed by hearing test. **b-e,** Effects on auditory function restoration. ABR **(b, d)** and DPOAE **(c, e)** thresholds in *Mir96^14C>A**/+**^* mice treated with AAV2-CMV-KKH-saCas9-sgRNA-4 ears (blue) and untreated ears (red) at 16 weeks age **(b-c)** and 26 weeks age **(d-e)**.

**Extended Data Fig. 6.**
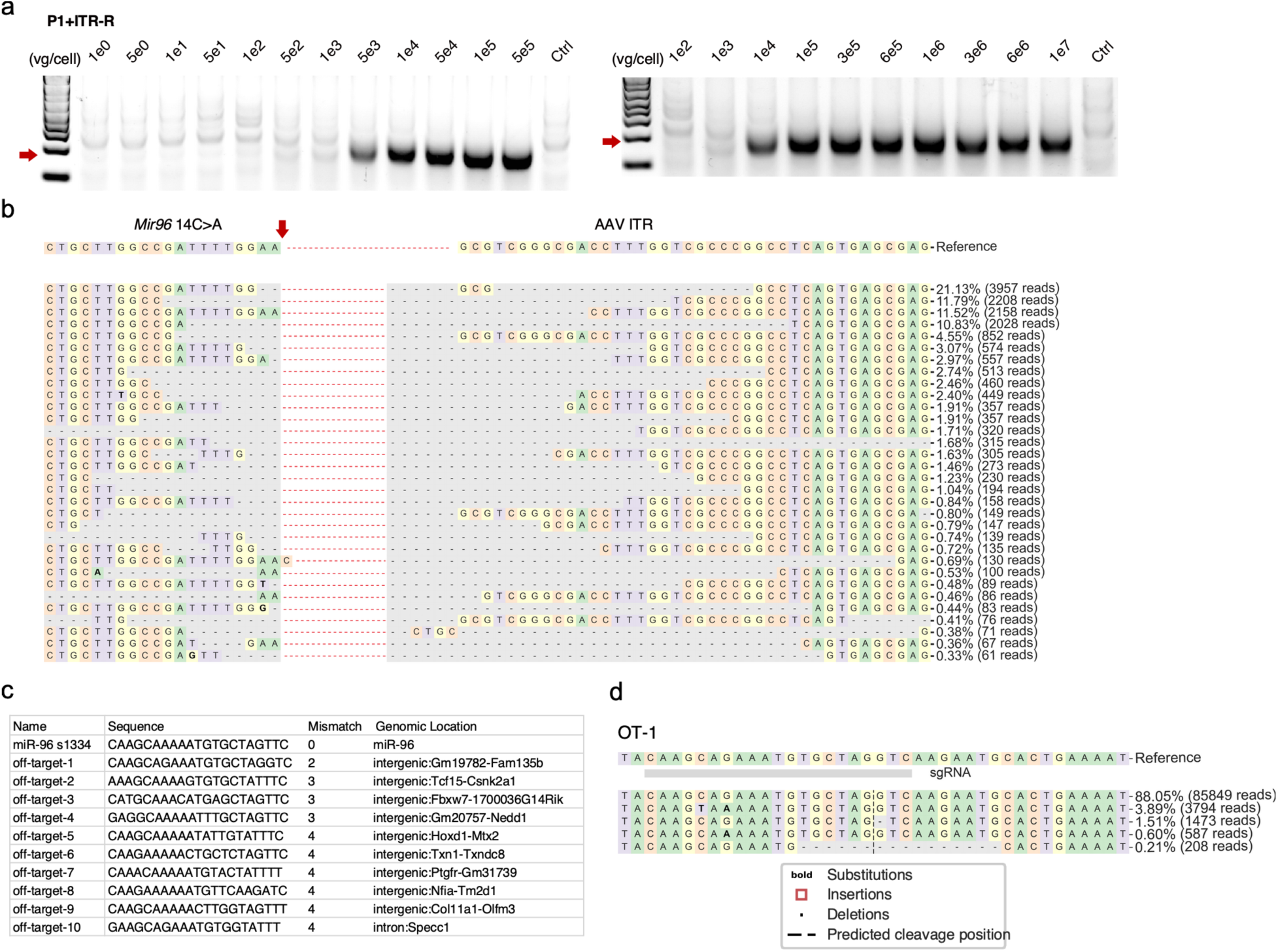
Safety assessment of AAV delivery of editing complex in adult mice. **a**, Gel image of the PCR showing the miR96-ITR integration fragment after different dosages of AAV treatment in HEK-miR96-14C>A cells. **b**, NGS results of miR96-ITR integration reads from AAV treated HEK-miR96-14C>A cells. **c**, The sequences of potential off-target genetic loci of KKH-saCas9/sgRNA-4 in mouse genome. None of these loci were associated with hearing function. **d**, The NGS result of off-target editing at the OT1 locus showed a low level of indel formation.

**Extended Data Fig. 7.**
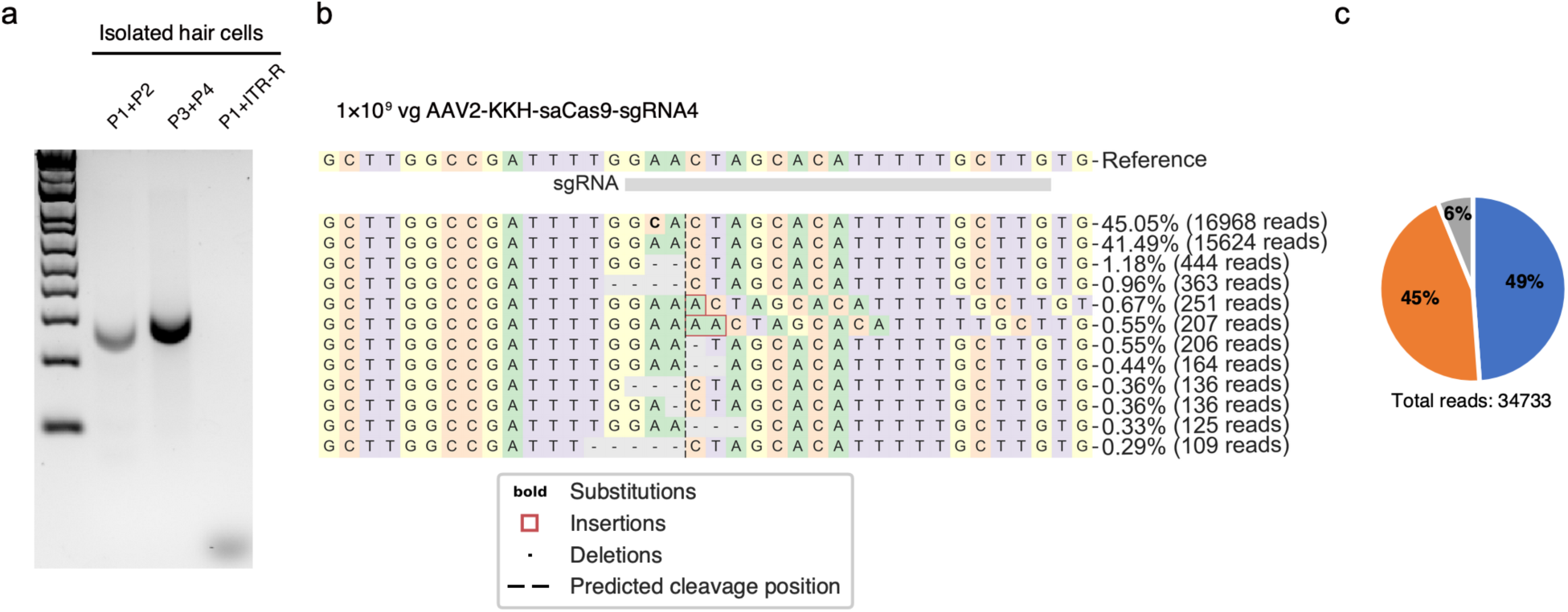
Low titer AAV2 eliminated AAV vector integration. **a**, Gel image of the PCR showing the KKH-saCas9 expression and miR96-ITR integration fragment after 1×10^9^ vg AAV2 treatment. **b, c**, Representative NGS results and pie chart analysis of 1×10^9^ vg AAV2-KKH-saCas9-sgRNA-4 injected ears from *Mir96^14C>A/+^* mice.

**Extended Data Fig. 8.**
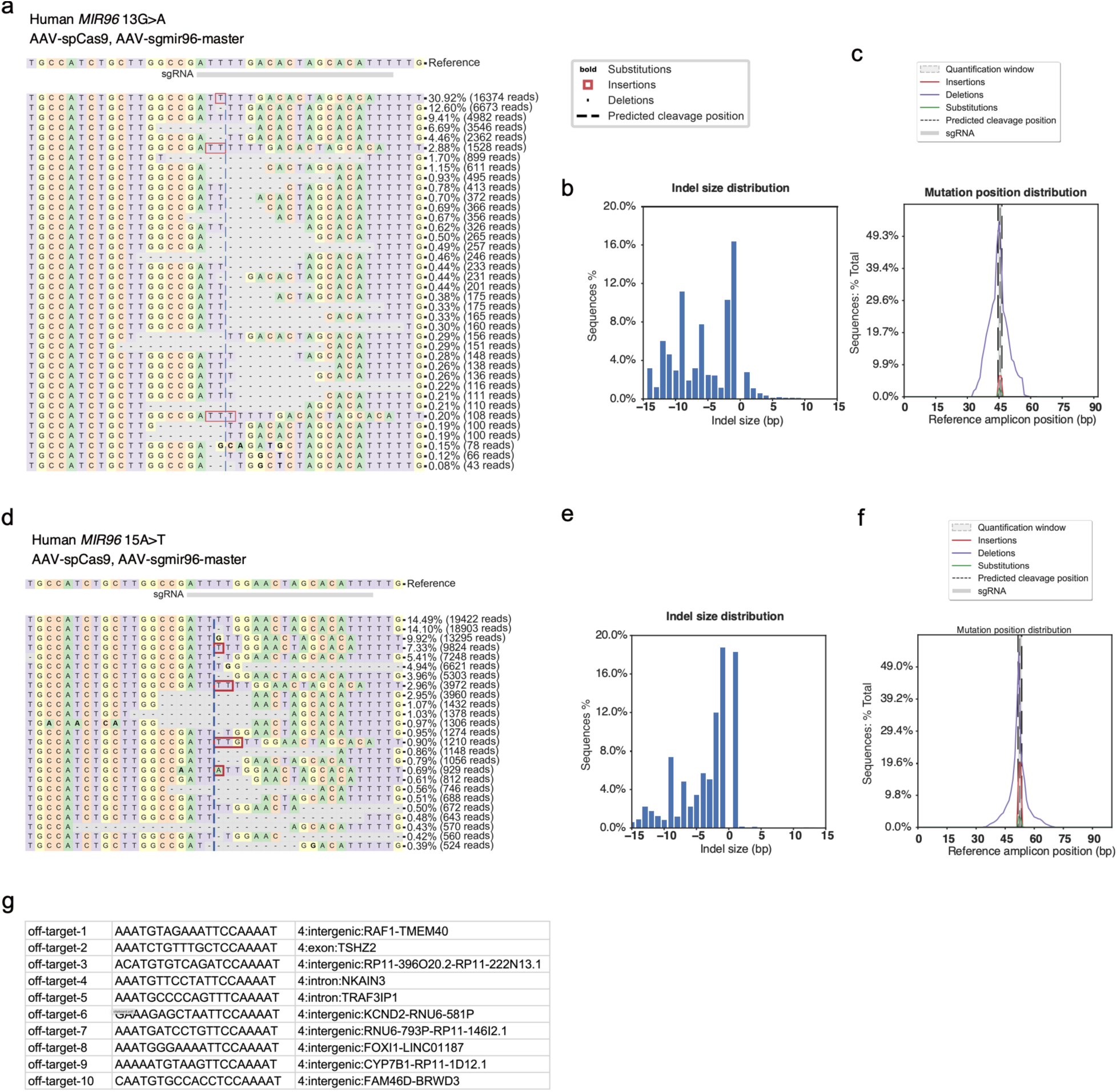
Efficient genome editing targeting multiple *MIR96* seed region mutations in human cells. **a,** Representative NGS results from spCas9/sgmiR96-master edited HEK-miR96 (13G to A) cells. **b, c,** Indel profiles from spCas9/sgmiR96-master edited HEK-miR96 (13G to A) cells. Minus numbers represent deletions, plus numbers represent insertions. **d,** Representative NGS results from spCas9/sgmiR96-master edited HEK-miR96 (14C to A) cells. **e, f,** Indel profiles from spCas9/sgmiR96-master edited HEK-miR96 (14C to A) cells. Minus numbers represent deletions, plus numbers represent insertions. **g,** The sequences of potential off-target genetic loci of spCas9/sgmiR96-master in human genome. None of these loci were associated with hearing function.

## Supplementary Information

**Supplementary Fig. 1.**
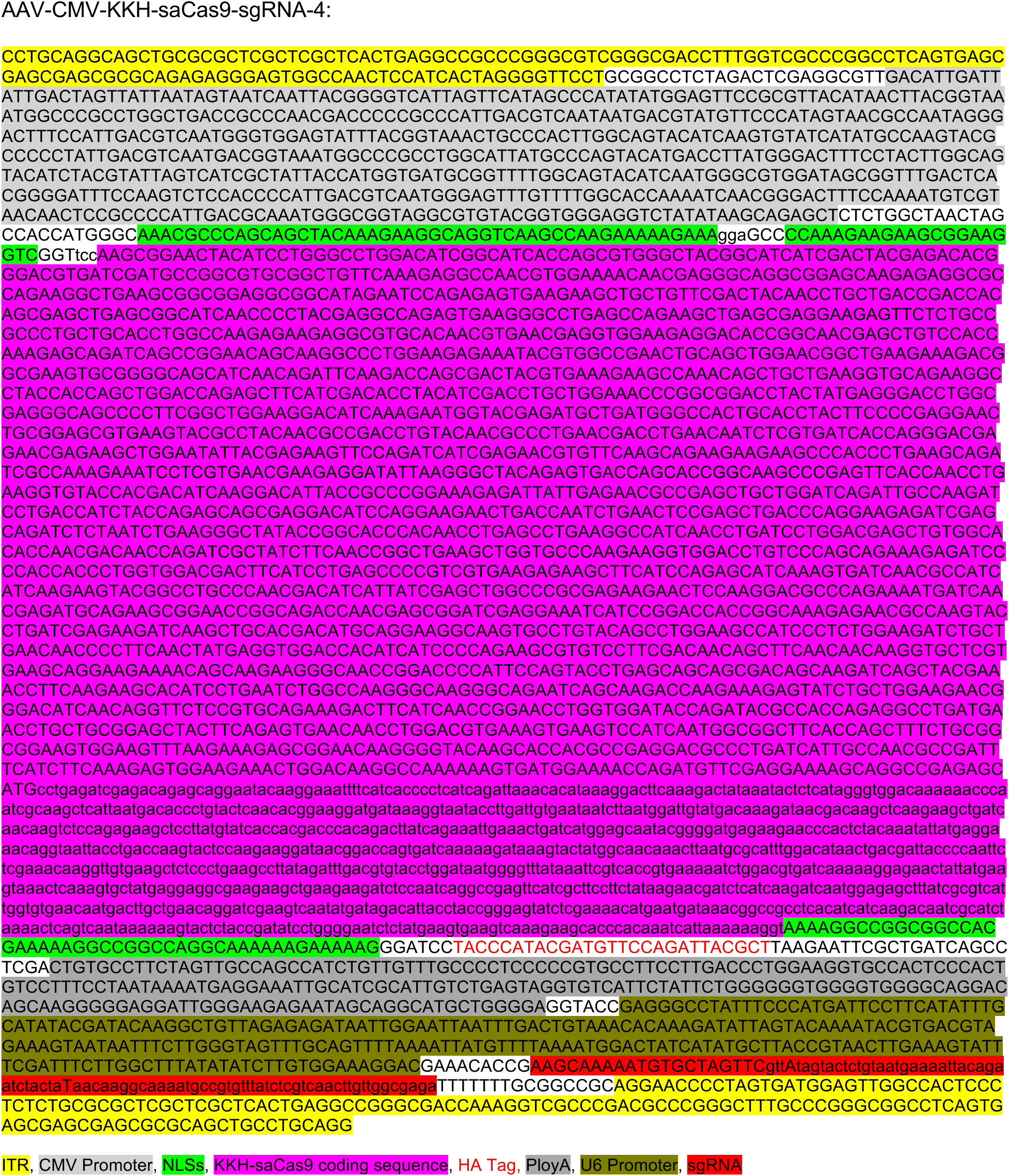
DNA sequence of AAV-CMV-KKH-saCas9-sgRNA-4.

**Supplementary Fig. 2.**
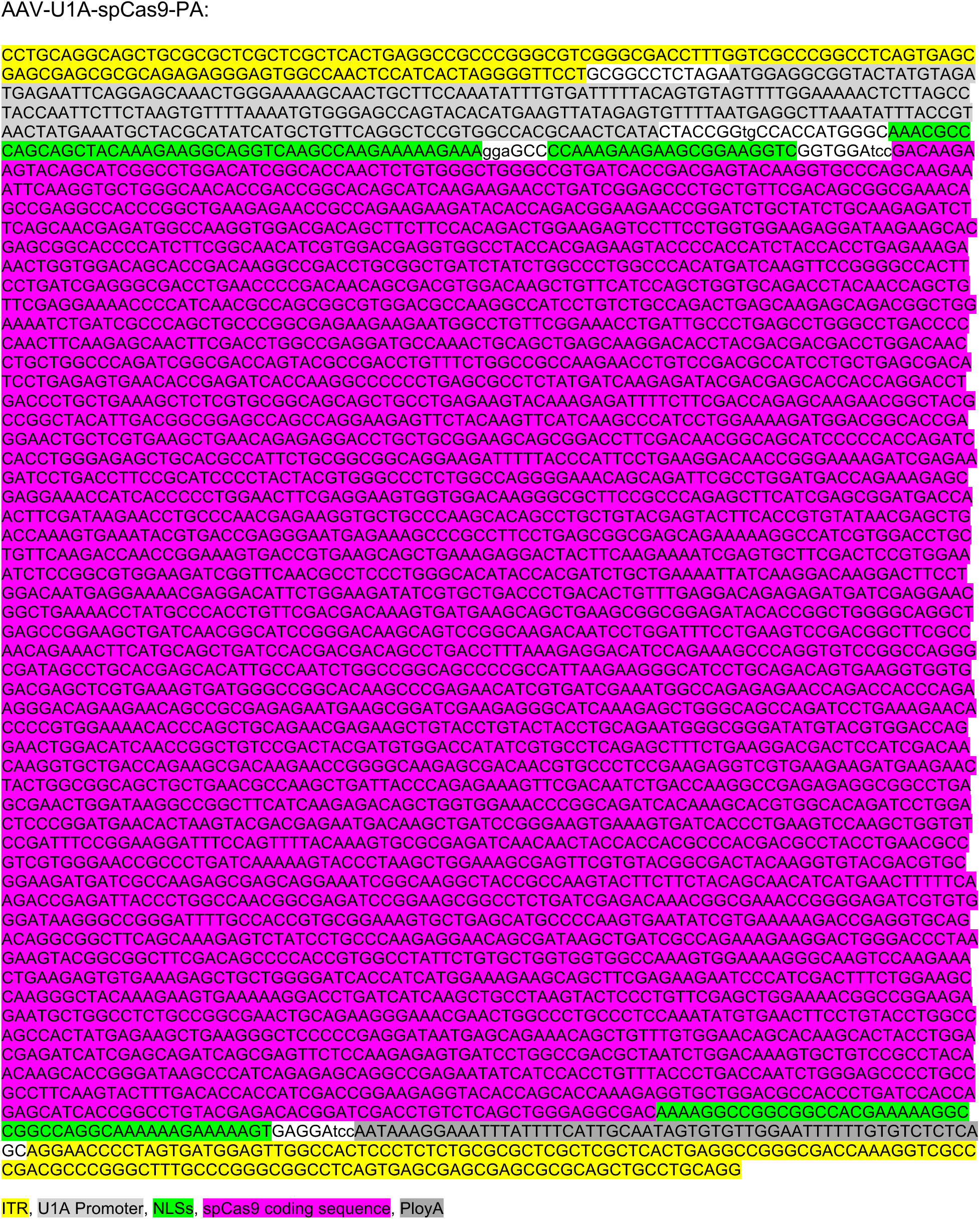
DNA sequence of AAV-U1A-spCas9-PA.

**Supplementary Fig. 3.**
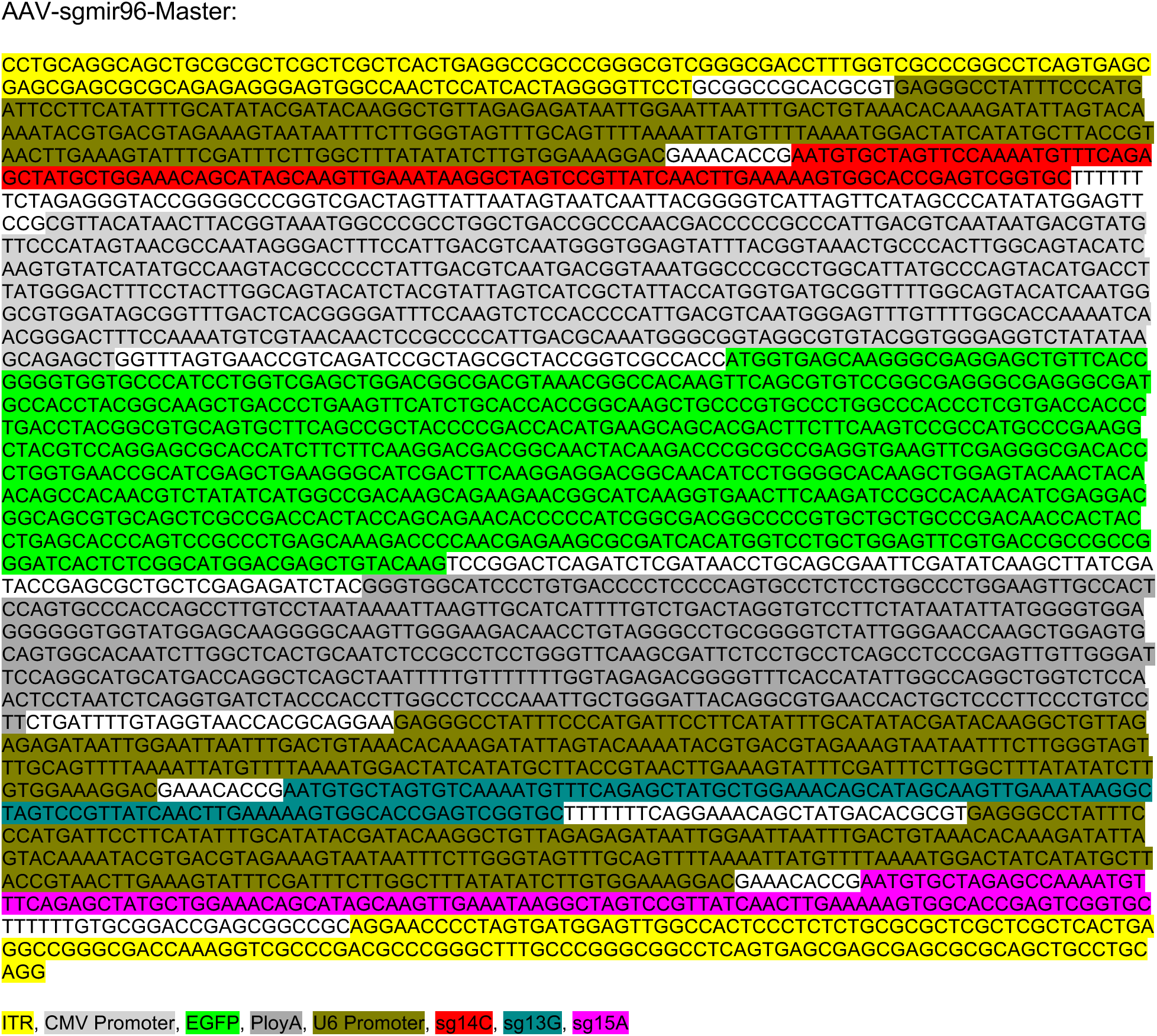
DNA sequence of AAV-sgmir96-Master.

**Supplementary Fig. 4.**
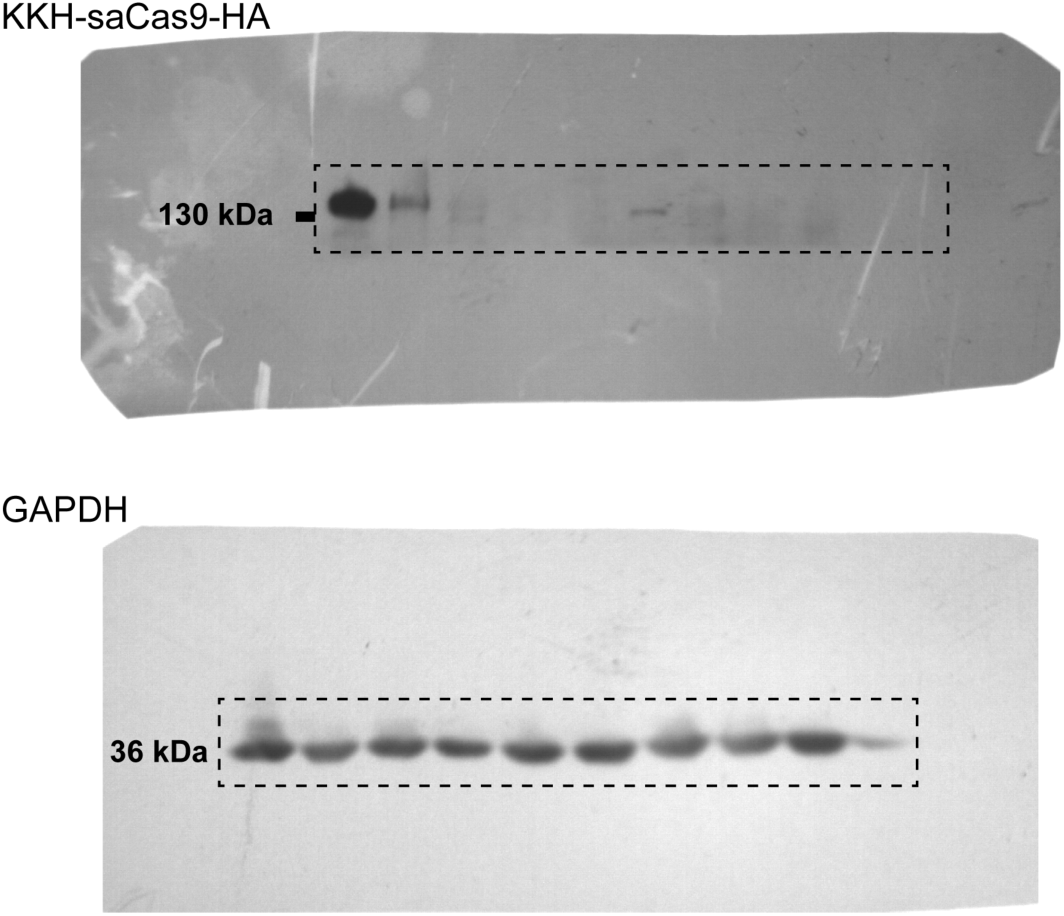
Uncropped images of western blots.

**Supplementary Table. 1.**
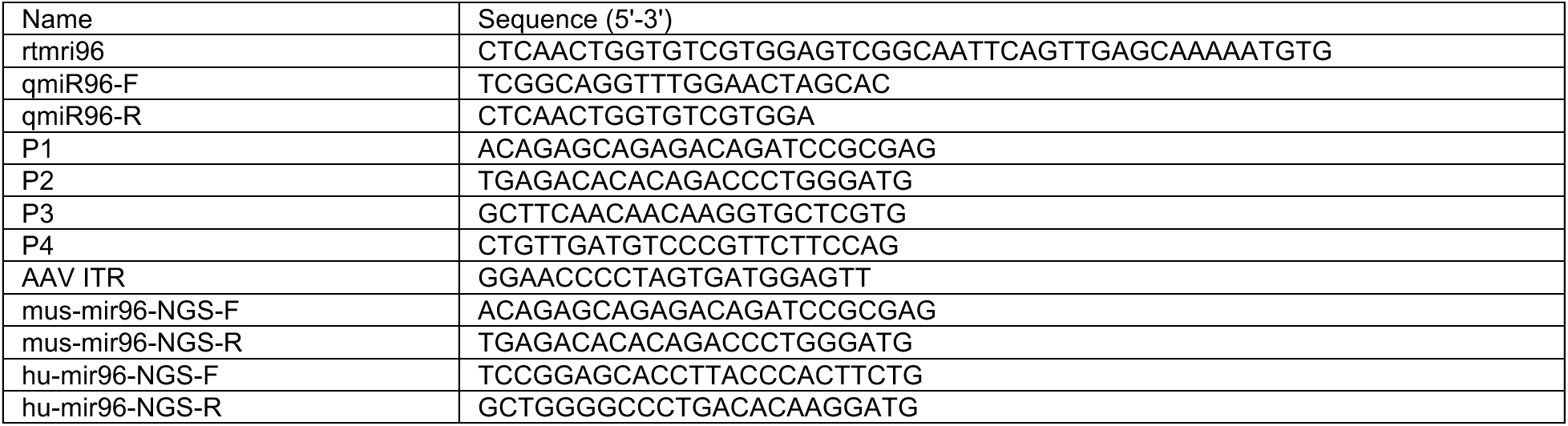
Primers used in this study.

**Supplementary Table. 2.**
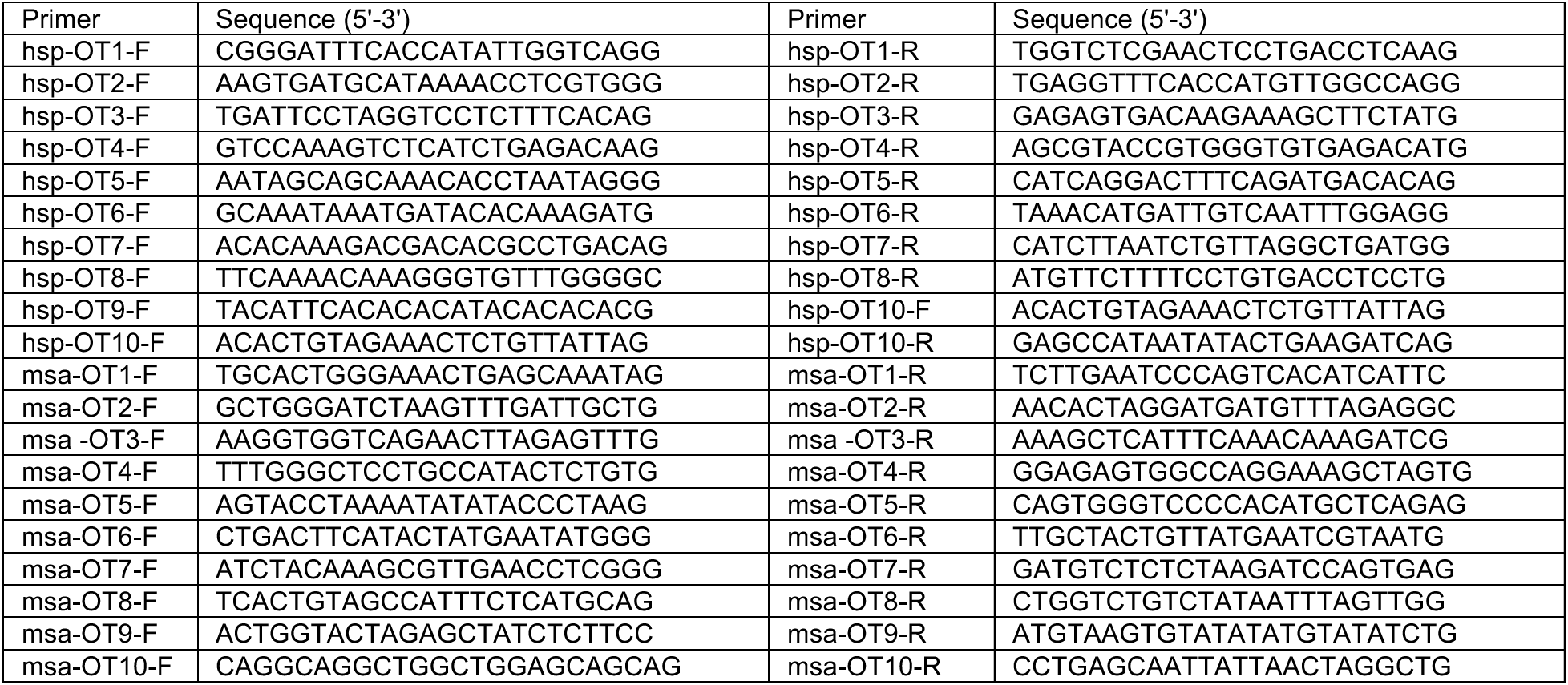
Primers used for offtarget analysis in this study.

**Supplementary Table. 3.**
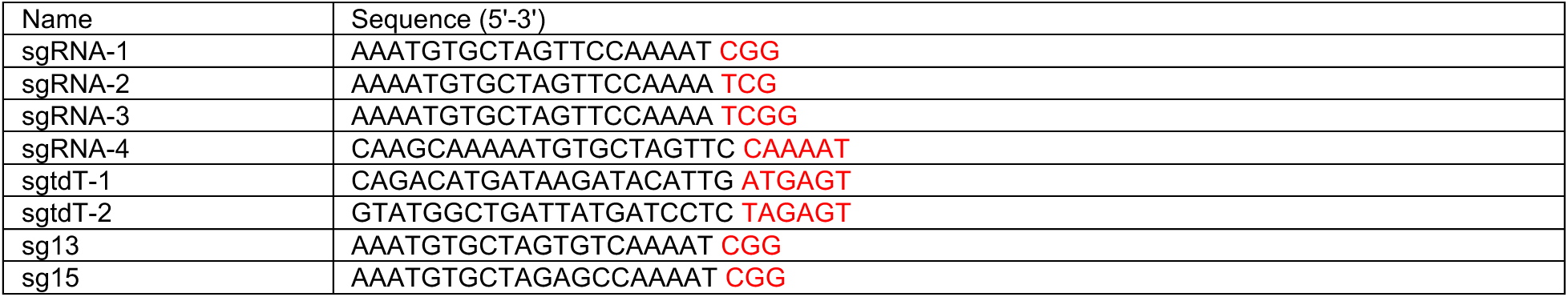
sgRNA protospacer sequences used in this study. Red fonts indicate the PAM sequence.

## References

1 Angeli, S., Lin, X. & Liu, X. Z. Genetics of hearing and deafness. Anat Rec (Hoboken) 295, 1812–1829, doi:10.1002/ar.22579 (2012).

2 Géléoc, G. S. & Holt, J. R. Sound strategies for hearing restoration. Science 344, 1241062, doi:10.1126/science.1241062 (2014).

3 Müller, U. & Barr-Gillespie, P. G. New treatment options for hearing loss. Nat Rev Drug Discov 14, 346–365, doi:10.1038/nrd4533 (2015).

4 Omichi, R., Shibata, S. B., Morton, C. C. & Smith, R. J. H. Gene therapy for hearing loss. Hum Mol Genet 28, R65–r79, doi:10.1093/hmg/ddz129 (2019).

5 Mahmoudian-Sani, M. R. et al. MicroRNAs: effective elements in ear-related diseases and hearing loss. Eur Arch Otorhinolaryngol 274, 2373–2380, doi:10.1007/s00405-017-4470-6 (2017).

6 Mendell, J. T. & Olson, E. N. MicroRNAs in stress signaling and human disease. Cell 148, 1172–1187, doi:10.1016/j.cell.2012.02.005 (2012).

7 Xiang, L. et al. miR-183/96 plays a pivotal regulatory role in mouse photoreceptor maturation and maintenance. Proc Natl Acad Sci U S A 114, 6376–6381, doi:10.1073/pnas.1618757114 (2017).

8 Kuhn, S. et al. miR-96 regulates the progression of differentiation in mammalian cochlear inner and outer hair cells. Proc Natl Acad Sci U S A 108, 2355–2360, doi:10.1073/pnas.1016646108 (2011).

9 Weston, M. D. et al. A mouse model of miR-96, miR-182 and miR-183 misexpression implicates miRNAs in cochlear cell fate and homeostasis. Sci Rep 8, 3569, doi:10.1038/s41598-018-21811-1 (2018).

10 Soldà, G. et al. A novel mutation within the MIR96 gene causes non-syndromic inherited hearing loss in an Italian family by altering pre-miRNA processing. Hum Mol Genet 21, 577–585, doi:10.1093/hmg/ddr493 (2012).

11 Mencía, A. et al. Mutations in the seed region of human miR-96 are responsible for nonsyndromic progressive hearing loss. Nat Genet 41, 609–613, doi:10.1038/ng.355 (2009).

12 Meola, N., Gennarino, V. A. & Banfi, S. microRNAs and genetic diseases. PathoGenetics 2, 7, doi:10.1186/1755-8417-2-7 (2009).

13 Weston, M. D., Pierce, M. L., Rocha-Sanchez, S., Beisel, K. W. & Soukup, G. A. MicroRNA gene expression in the mouse inner ear. Brain Res 1111, 95–104, doi:10.1016/j.brainres.2006.07.006 (2006).

14 Schlüter, T. et al. miR-96 is required for normal development of the auditory hindbrain. Hum Mol Genet 27, 860–874, doi:10.1093/hmg/ddy007 (2018).

15 Xu, S., Witmer, P. D., Lumayag, S., Kovacs, B. & Valle, D. MicroRNA (miRNA) transcriptome of mouse retina and identification of a sensory organ-specific miRNA cluster. J Biol Chem 282, 25053–25066, doi:10.1074/jbc.M700501200 (2007).

16 Wienholds, E. et al. MicroRNA expression in zebrafish embryonic development. Science 309, 310–311, doi:10.1126/science.1114519 (2005).

17 Lewis, M. A. et al. An ENU-induced mutation of miR-96 associated with progressive hearing loss in mice. Nat Genet 41, 614–618, doi:10.1038/ng.369 (2009).

18 Lewis, M. A., Di Domenico, F., Ingham, N. J., Prosser, H. M. & Steel, K. P. Hearing impairment due to Mir183/96/182 mutations suggests both loss and gain of function effects. Dis Model Mech 14, doi:10.1242/dmm.047225 (2020).

19 Gao, X. et al. Treatment of autosomal dominant hearing loss by in vivo delivery of genome editing agents. Nature 553, 217–221, doi:10.1038/nature25164 (2018).

20 Tao, Y. et al. Treatment of monogenic and digenic dominant genetic hearing loss by CRISPR-Cas9 ribonucleoprotein delivery in vivo. Nat Commun 14, 4928, doi:10.1038/s41467-023-40476-7 (2023).

21 György, B. et al. Allele-specific gene editing prevents deafness in a model of dominant progressive hearing loss. Nat Med 25, 1123–1130, doi:10.1038/s41591-019-0500-9 (2019).

22 Wu, J. et al. Single and Dual Vector Gene Therapy with AAV9-PHP.B Rescues Hearing in Tmc1 Mutant Mice. Mol Ther 29, 973–988, doi:10.1016/j.ymthe.2020.11.016 (2021).

23 Gu, X. et al. Prevention of acquired sensorineural hearing loss in mice by in vivo Htra2 gene editing. Genome Biol 22, 86, doi:10.1186/s13059-021-02311-4 (2021).

24 Xue, Y. et al. Gene editing in a Myo6 semi-dominant mouse model rescues auditory function. Mol Ther 30, 105–118, doi:10.1016/j.ymthe.2021.06.015 (2022).

25 Noh, B. et al. In vivo outer hair cell gene editing ameliorates progressive hearing loss in dominant-negative Kcnq4 murine model. Theranostics 12, 2465–2482, doi:10.7150/thno.67781 (2022).

26 Cui, C. et al. Precise detection of CRISPR-Cas9 editing in hair cells in the treatment of autosomal dominant hearing loss. Mol Ther Nucleic Acids 29, 400–412, doi:10.1016/j.omtn.2022.07.016 (2022).

27 Yeh, W. H. et al. In vivo base editing restores sensory transduction and transiently improves auditory function in a mouse model of recessive deafness. Sci Transl Med 12, doi:10.1126/scitranslmed.aay9101 (2020).

28 Jiang, L., Wang, D., He, Y. & Shu, Y. Advances in gene therapy hold promise for treating hereditary hearing loss. Mol Ther 31, 934–950, doi:10.1016/j.ymthe.2023.02.001 (2023).

29 Yoshimura, H., Shibata, S. B., Ranum, P. T., Moteki, H. & Smith, R. J. H. Targeted Allele Suppression Prevents Progressive Hearing Loss in the Mature Murine Model of Human TMC1 Deafness. Mol Ther 27, 681–690, doi:10.1016/j.ymthe.2018.12.014 (2019).

30 Morag A. Lewis, M. L., Miguel Angel Moreno-Pelayo and Karen P. Steel. in Association for Research in Otolaryngology midwinter meeting abstracts (2020).

31 Jinek, M. et al. A programmable dual-RNA-guided DNA endonuclease in adaptive bacterial immunity. Science 337, 816–821, doi:10.1126/science.1225829 (2012).

32 Chatterjee, P. et al. An engineered ScCas9 with broad PAM range and high specificity and activity. Nat Biotechnol 38, 1154–1158, doi:10.1038/s41587-020-0517-0 (2020).

33 Kleinstiver, B. P. et al. Broadening the targeting range of Staphylococcus aureus CRISPR-Cas9 by modifying PAM recognition. Nat. Biotechnol. 33, 1293–1298, doi:10.1038/nbt.3404 (2015).

34 Hu, Z. et al. A compact Cas9 ortholog from Staphylococcus Auricularis (SauriCas9) expands the DNA targeting scope. PLoS Biol 18, e3000686, doi:10.1371/journal.pbio.3000686 (2020).

35 Schmid-Burgk, J. L. et al. Highly Parallel Profiling of Cas9 Variant Specificity. Mol Cell 78, 794–800.e798, doi:10.1016/j.molcel.2020.02.023 (2020).

36 Anzalone, A. V. et al. Search-and-replace genome editing without double-strand breaks or donor DNA. Nature 576, 149–157, doi:10.1038/s41586-019-1711-4 (2019).

37 Chen, P. J. & Liu, D. R. Prime editing for precise and highly versatile genome manipulation. Nat Rev Genet 24, 161–177, doi:10.1038/s41576-022-00541-1 (2023).

38 Nelson, J. W. et al. Engineered pegRNAs improve prime editing efficiency. Nat Biotechnol 40, 402–410, doi:10.1038/s41587-021-01039-7 (2022).

39 Omichi, R., Yoshimura, H., Shibata, S. B., Vandenberghe, L. H. & Smith, R. J. H. Hair Cell Transduction Efficiency of Single- and Dual-AAV Serotypes in Adult Murine Cochleae. Mol Ther Methods Clin Dev 17, 1167–1177, doi:10.1016/j.omtm.2020.05.007 (2020).

40 Du, W. et al. Rescue of auditory function by a single administration of AAV-TMPRSS3 gene therapy in aged mice of human recessive deafness DFNB8. Mol Ther, doi:10.1016/j.ymthe.2023.05.005 (2023).

41 Wu, Y. et al. Highly efficient therapeutic gene editing of human hematopoietic stem cells. Nat Med 25, 776–783, doi:10.1038/s41591-019-0401-y (2019).

42 Koblan, L. W. et al. Improving cytidine and adenine base editors by expression optimization and ancestral reconstruction. Nat Biotechnol 36, 843–846, doi:10.1038/nbt.4172 (2018).

43 Chen, B. et al. Expanding the CRISPR imaging toolset with Staphylococcus aureus Cas9 for simultaneous imaging of multiple genomic loci. Nucleic Acids Res 44, e75, doi:10.1093/nar/gkv1533 (2016).

44 Staahl, B. T. et al. Efficient genome editing in the mouse brain by local delivery of engineered Cas9 ribonucleoprotein complexes. Nat Biotechnol 35, 431–434, doi:10.1038/nbt.3806 (2017).

45 Meyers, J. R. et al. Lighting up the senses: FM1-43 loading of sensory cells through nonselective ion channels. J Neurosci 23, 4054–4065, doi:10.1523/jneurosci.23-10-04054.2003 (2003).

46 Shu, Y. et al. Renewed proliferation in adult mouse cochlea and regeneration of hair cells. Nat Commun 10, 5530, doi:10.1038/s41467-019-13157-7 (2019).

47 Brooks, A. R. et al. Transcriptional silencing is associated with extensive methylation of the CMV promoter following adenoviral gene delivery to muscle. J Gene Med 6, 395–404, doi:10.1002/jgm.516 (2004).

48 Levy, J. M. et al. Cytosine and adenine base editing of the brain, liver, retina, heart and skeletal muscle of mice via adeno-associated viruses. Nat Biomed Eng 4, 97–110, doi:10.1038/s41551-019-0501-5 (2020).

49 Gray, S. J. et al. Optimizing promoters for recombinant adeno-associated virus-mediated gene expression in the peripheral and central nervous system using self-complementary vectors. Hum Gene Ther 22, 1143–1153, doi:10.1089/hum.2010.245 (2011).

50 Hanlon, K. S. et al. High levels of AAV vector integration into CRISPR-induced DNA breaks. Nat Commun 10, 4439, doi:10.1038/s41467-019-12449-2 (2019).

51 Nelson, C. E. et al. Long-term evaluation of AAV-CRISPR genome editing for Duchenne muscular dystrophy. Nat Med 25, 427–432, doi:10.1038/s41591-019-0344-3 (2019).

52 Tsai, S. Q. et al. CIRCLE-seq: a highly sensitive in vitro screen for genome-wide CRISPR-Cas9 nuclease off-targets. Nat Methods 14, 607–614, doi:10.1038/nmeth.4278 (2017).

53 Doench, J. G. et al. Optimized sgRNA design to maximize activity and minimize off-target effects of CRISPR-Cas9. Nat Biotechnol 34, 184–191, doi:10.1038/nbt.3437 (2016).

54 Tycko, J. et al. Pairwise library screen systematically interrogates Staphylococcus aureus Cas9 specificity in human cells. Nat Commun 9, 2962, doi:10.1038/s41467-018-05391-2 (2018).

55 Rudnicki, A. & Avraham, K. B. microRNAs: the art of silencing in the ear. EMBO Mol Med 4, 849–859, doi:10.1002/emmm.201100922 (2012).

56 Li, H., Kloosterman, W. & Fekete, D. M. MicroRNA-183 family members regulate sensorineural fates in the inner ear. J Neurosci 30, 3254–3263, doi:10.1523/jneurosci.4948-09.2010 (2010).

57 Xiong, H. et al. Modulation of miR-34a/SIRT1 signaling protects cochlear hair cells against oxidative stress and delays age-related hearing loss through coordinated regulation of mitophagy and mitochondrial biogenesis. Neurobiol Aging 79, 30–42, doi:10.1016/j.neurobiolaging.2019.03.013 (2019).

58 Miguel, V. et al. The Role of MicroRNAs in Environmental Risk Factors, Noise-Induced Hearing Loss, and Mental Stress. Antioxidants & Redox Signaling 28, 773–796, doi:10.1089/ars.2017.7175 (2017).

59 Tan, P. X. et al. MicroRNA-207 enhances radiation-induced apoptosis by directly targeting Akt3 in cochlea hair cells. Cell Death Dis 5, e1433, doi:10.1038/cddis.2014.407 (2014).

60 Kuzmin, D. A. et al. The clinical landscape for AAV gene therapies. Nat Rev Drug Discov 20, 173–174, doi:10.1038/d41573-021-00017-7 (2021).

61 Wang, D., Zhang, F. & Gao, G. CRISPR-Based Therapeutic Genome Editing: Strategies and In Vivo Delivery by AAV Vectors. Cell 181, 136–150, doi:10.1016/j.cell.2020.03.023 (2020).

62 Yuen, C. T. L. et al. High-fidelity KKH variant of Staphylococcus aureus Cas9 nucleases with improved base mismatch discrimination. Nucleic Acids Res 50, 1650–1660, doi:10.1093/nar/gkab1291 (2022).

63 Akcakaya, P. et al. In vivo CRISPR editing with no detectable genome-wide off-target mutations. Nature 561, 416–419, doi:10.1038/s41586-018-0500-9 (2018).

64 Davis, K. M., Pattanayak, V., Thompson, D. B., Zuris, J. A. & Liu, D. R. Small molecule-triggered Cas9 protein with improved genome-editing specificity. Nat Chem Biol 11, 316–318, doi:10.1038/nchembio.1793 (2015).

65 McCown, T. J., Xiao, X., Li, J., Breese, G. R. & Samulski, R. J. Differential and persistent expression patterns of CNS gene transfer by an adeno-associated virus (AAV) vector. Brain Res 713, 99–107, doi:10.1016/0006-8993(95)01488-8 (1996).

66 Paterna, J. C., Moccetti, T., Mura, A., Feldon, J. & Büeler, H. Influence of promoter and WHV post-transcriptional regulatory element on AAV-mediated transgene expression in the rat brain. Gene Ther 7, 1304–1311, doi:10.1038/sj.gt.3301221 (2000).

67 Zhu, W. et al. Precisely controlling endogenous protein dosage in hPSCs and derivatives to model FOXG1 syndrome. Nat Commun 10, 928, doi:10.1038/s41467-019-08841-7 (2019).

68 Ran, F. A. et al. Genome engineering using the CRISPR-Cas9 system. Nat. Protoc. 8, 2281–2308, doi:10.1038/nprot.2013.143 (2013).

69 Ran, F. A. et al. In vivo genome editing using Staphylococcus aureus Cas9. Nature 520, 186–191, doi:10.1038/nature14299 (2015).

70 Wang, D. et al. Cas9-mediated allelic exchange repairs compound heterozygous recessive mutations in mice. Nat Biotechnol 36, 839–842, doi:10.1038/nbt.4219 (2018).

71 Duan, Y. et al. The Clustered, Regularly Interspaced, Short Palindromic Repeats-associated Endonuclease 9 (CRISPR/Cas9)-created MDM2 T309G Mutation Enhances Vitreous-induced Expression of MDM2 and Proliferation and Survival of Cells. J Biol Chem 291, 16339–16347, doi:10.1074/jbc.M116.729467 (2016).

72 Clement, K. et al. CRISPResso2 provides accurate and rapid genome editing sequence analysis. Nat Biotechnol 37, 224–226, doi:10.1038/s41587-019-0032-3 (2019).

